# Environmental DNA is more effective than hand sorting in evaluating earthworm biodiversity recovery under regenerative agriculture

**DOI:** 10.1101/2023.11.07.565975

**Authors:** J. Llanos, H. Hipperson, G. Horsburgh, M.G. Lappage, K.H. Maher, T. Burke, J.R. Leake, P. J. Watt

## Abstract

Regenerating soil biodiversity is vital to help reverse declines in the health of agricultural soils caused by intensification, and to support sustainable food production and agro-ecosystem services. Earthworms are key functional components of soil biodiversity, with different ecotypes and species delivering specific beneficial soil functions. However, conventional monitoring by hand-sorting from soil pits is highly labour intensive, can reliably identify only adults to species, and may under-record anecics (deep-burrowing ecotypes). Here, we compare soil environmental DNA (eDNA) metabarcoding using two different primer sets and next-generation sequencing, with hand-sorting from standard soil-pits. The experiment comprised four conventionally managed arable fields into which strips of grass-clover ley had been introduced three years earlier. Earthworm population responses had been recorded by hand-sorting for the first two years and our goal was to assess these in the third year and to compare them both by hand and with the use of eDNA. The eDNA method found the same 8 species as hand-sorting, but had greater power for detecting anecic earthworms and quantifying local species richness. Earthworm abundance increased by over 55% into the third year of the leys, surpassing abundances in adjacent permanent grasslands, helping to explain the observed soil health regeneration. Both overall relative abundances and site occupancy proportions of earthworm eDNA were found to have potential as proxies for abundance, and the performance of each of these measures and the implications for further work are discussed. We demonstrate that eDNA can overcome some of the significant barriers and limitations to monitoring earthworm diversity and recommend its wider use both to better understand the earthworm population benefits of different soil management practices, and to guide agricultural policy and practice decisions affecting soil health.

## 1. Introduction

It has been estimated that humanity derives 98.8% of its food calories from soils (Kopittke et al., 2019), and soils temporarily store and filter most of the fresh water used domestically and in agriculture, so protecting and enhancing this resource is essential for sustaining the predicted 9.7 billion people on the planet in 2050 (UN, 2019). Intensive cropping using annual ploughing and harrowing depletes functional biodiversity, including earthworm populations (Edwards and Lofty 1982; Briones and Schmidt 2017), and destroys soil aggregates in which carbon, nutrients and water are stored (Low, 1972; Guest et al., 2022). This leads to the loss of macropores, which impairs drainage, and increases the risk of crop failure, flooding, erosion, and water pollution (Holden et al., 2019; Berdeni et al., 2021). Currently, rates of soil erosion often far outstrip rates of soil formation in conventional arable cropping using intensive tillage, frequently reducing the lifespans of topsoils (to 30 cm depth) to less than 200 years (Evans et al., 2020). For England and Wales, the economic loss from soil degradation has been calculated at *ca* £1.2 billion per year (Graves et al., 2015). Of this loss, 47% is attributable to depletion of organic matter, mainly from intensively cultivated soils, which limits their capacity to store water and nutrients and reduces the food supply and habitat quality for most heterotrophic soil organisms, including earthworms. A further 39% of the estimated economic loss is due to soil compaction, which is exacerbated by weak soil structure due to the loss of water-stable macroaggregates and the macropores that are generated, especially by plant roots and earthworms (Tisdall and Oades, 1982; Hallam and Hodson, 2020; Hallam et al., 2020). In turn, this causes poor soil drainage, contributing to erosion, which accounts for a further 12% of the economic loss (Graves et al., 2015).

Conservation tillage practices that reduce soil disturbance and keep it covered can improve soil aggregation, carbon sequestration and earthworm abundance (Briones & Schmidt 2017; Giannitsopoulos et al., 2019), and extend topsoil lifespans to over 10,000 years (Evans et al., 2020). Farmers are increasingly aware of the need to manage soils more sustainably, with rapid adoption of conservation agriculture (Kassam et al., 2019) and regenerative agriculture practices (Jaworski et al., 2023) to improve soil health and functional biodiversity. These approaches aim to minimise soil disturbance, diversify rotations, keep soil covered with crop residues (FAO, 2023), and feed soil organisms via living roots using cover crops and leys (Jaworski et al., 2023). Recent policy developments, such as England’s Environmental Improvement Plan and Sustainable Farming Incentive payments (DEFRA, 2023), the Sustainable Farming Scheme in Wales and the upcoming EU Soil Health Law (European Commission, 2022), aim to promote adoption of these kinds of more sustainable soil management and to improve soil health. Guiding best practice for sustainable soil management requires efficient and effective tools and indicators to assess soil health and quality (Karlen et al., 2019; Liu et al., 2020; Lehmann et al., 2020). Healthy soils are characterised by chemical, physical and biological attributes that interact to deliver functions and ecosystem services (Lehmann et al., 2020). These include crop and fodder production, nutrient fertility, water-storage and filtration capacity, carbon storage and sequestration, and biotic communities that help control pests and diseases and improve soil aggregation and porosity, and liberate mineral nutrients to crops (Rinot et al., 2019; Giannitsopoulos et al., 2019; Lehmann et al., 2020; Austen et al., 2022; Guest et al., 2022).

Earthworms are widely recognised as bio-indicators of healthy soils by farmers and scientists (Stroud et al., 2019; Lehmann et al., 2020; Hallam et al., 2021). They are ‘ecosystem engineers’ because of their large effects on the structure and functioning of soils (Lavelle et al., 1997; Cunha et al., 2016). They have been categorised into three ecotypes based on their soil occupancy and feeding behaviours, with epigeic types living near the surface and feeding on plant litter, endogeic types burrowing through and feeding within the topsoil, and anecic types constructing deep vertical burrows at the top of which they gather plant litter into middens (Stroud & Goulding, 2022). As a result of their burrowing and casting, earthworms create macropores and macroaggregates (Hallam and Hodson, 2020; Hallam et al., 2020) in which organic carbon is sequestered and protected from microbial degradation, whilst at the same time stimulating soil microbial activities (Zhang et al., 2013). Their generation of macropores and macroaggregates increases soil water-holding capacity, gas exchange and drainage (Zhang and Schrader, 1993; Blanchart et al., 1999; Schaik et al., 2014; Sheehy et al., 2019; Hallam and Hodson, 2020). Via their feeding, earthworms are responsible for accelerating the breakdown of organic matter and distributing it deeper into the soil profile, whilst increasing the availability of mineral nutrients that improve crop yields and microbial activity (Lavelle et al., 1998; Chaoui et al., 2003; Le Bayon and Milleret, 2009). The biological properties of soils are also greatly affected by earthworms because of their direct or indirect interactions with other organisms, and their ability to stimulate microbial activity (Binet et al., 1998; Monroy et al., 2011; Bart et al., 2019; Wang et al., 2020). Preferential burial of mycotoxin-producing *Fusarium*-infecting wheat straw by specific deep-burrowing ecotypes like *Lumbricus terrestris* can help to reduce phytopathogenic and toxinogenic fungal inoculum in conservation agriculture where straw is not ploughed in (Wolfarth et al., 2011). As well as maintaining and promoting agricultural soil health, a meta-analysis by van Groenigen et al. (2014) concluded that earthworms increased crop yields by an average of 25%, with higher increases seen in nitrogen-deficient soils or those that had previously been disturbed and lost their structure.

Despite the immense contribution of earthworms to soil health in agroecosystems, the lack of efficient tools to monitor their populations constrains their routine use as soil health indicators, with their populations understudied and poorly mapped (e.g. Earthworm Society of Britain 2020). There has been some recent progress in the broad-scale mapping of distributions and diversity nationally (e.g. UK, Ashwood et al. 2024) and internationally (Phillips et al., 2019) but the tools available to farmers are time consuming, primitive and constrained by a lack of taxonomic expertise (Jiménez et al., 2006; Čoja et al., 2008; Bartlett et al., 2010; Andriuzzi et al., 2017; Stroud and Goulding, 2019, 2022). In sum, our knowledge of earthworm distributions and diversity remains extremely limited.

Conventional monitoring of earthworm populations is challenging because they are seasonally dynamic, strongly affected by weather (especially dry periods prior to sampling), and populations are normally dominated by juveniles that can only reliably be identified to ecotype (Prendergast-Miller et al., 2021), as standard keys use adult characteristics (Sherlock 2018). Knowledge of species abundance, and not just total earthworm numbers or ecotypes, is needed if we are to better understand their distributions and biodiversity responses to farmland management, as there are important functional differences between species within an ecotype (Hoeffner et al., 2022). Addressing these limitations, and developing more standardised methods for sampling earthworm populations and species, are therefore vital for monitoring their distributions and realising their potential as indicators and promoters of soil health.

Environmental DNA (eDNA) sampling is a promising alternative method for assessing earthworm populations that could address many of the limitations of traditional methods. eDNA sampling involves collecting, amplifying and sequencing genetic material that has been left behind by organisms in their environment, and which may originate from deposits such as hair, skin cells, mucus or faeces (Belle et al., 2019). For species like earthworms, with spatially and seasonally highly variable populations, eDNA offers potential advantages over manual extraction and hand sorting from soil pits. Most studies employing eDNA sampling have tended to focus on aquatic environments (Belle et al., 2019), but the feasibility of applying eDNA sampling techniques to terrestrial environments has also been demonstrated. This includes sampling of ice cores, sediments, plant material, scats and air (for example Hofreiter et al., 2003; Willerslev et al., 2007; Bohmann et al., 2011; Thomsen and Sigsgaard, 2019; Bohmann and Lynggaard, 2023).

Soil is a reservoir of eDNA that can be sampled to build a picture of both above- and below-ground biodiversity. For example, Yoccoz et al. (2012) showed that soil eDNA profiles were consistent with above-ground plant diversity, and Andersen et al. (2012) sampled soil with known species compositions to demonstrate the viability of soil eDNA for vertebrate biodiversity sampling. Despite their importance for soil health, both soil meso- and macrofauna are underrepresented in agricultural eDNA monitoring studies, with just 7% of publications to date describing work in this area and the vast majority (93%) focused on soil microbiota (Kestel et. al., 2022). Bienert et al. (2012) were the first to demonstrate that eDNA can be used for sampling earthworm communities in undisturbed woodland and meadow soils. This was followed by Pansu et al. (2015), who used eDNA to show differences in earthworm diversity at the landscape level in the northern French Alps. To date, earthworm eDNA sampling of arable soils has received virtually no attention, except for a recent study in Denmark, which compared hand sorting from 36 soil pits with eDNA in 18 small field plots of spring barley, comprising control, pig slurry and mineral fertiliser, replicated three times on two soil types (Lilja et al., 2023). They found five earthworm species by hand sorting and eight by eDNA, but whilst hand sorting revealed significant effects of soil type and fertiliser treatments, these were not detected by eDNA based on 0.25–10 g soil samples.

An important priority for future research of this kind is validating the use of eDNA in studies of earthworm biodiversity and population responses to change from conventional arable cropping to regenerative agricultural practices that aim to enable recovery of beneficial organisms that improve soil health (Jaworski et al., 2023). Amongst the most effective regenerative agriculture practices is the reintroduction of grass-clover leys into arable rotations, which promotes rapid recovery of earthworm populations in 1–2-year leys (Prendergast-Miller et al., 2021), and regenerates soil aggregates (Guest et al., 2022) and soil fertility (Austen et al., 2022).

Here, we assess the performance of eDNA analysis in comparison with hand-sorting for earthworm biodiversity and abundance, in four long-term arable fields containing 3-year grass-clover leys, taking advantage of the existing ‘SoilBioHedge’ experiment (Hallam et al., 2020). SoilBioHedge has provided detailed information on earthworm populations in these arable fields, 1–2-year ley strips, grassy margins and hedgerows, and in the adjoining permanent grasslands (Holden et al., 2019, Prendergast-Miller et al., 2021). It has also reported the effect of earthworms on soil properties including hydrological functioning from soil monolith studies in the same arable fields (Hallam et al., 2020), and how the grass-clover leys improve soil hydrology, soil structure, carbon storage and crop performance (Berdeni et al., 2021; Guest et al., 2022). However, SoilBioHedge did not assess the effects of a third year of grass-clover ley on earthworm populations, which has important implications for decision-making about the optimal duration of leys in regenerative agriculture.

Soil samples for earthworm eDNA were spatially paired with standardised soil-pits and earthworm hand-sorting and counting, with adults identified to species-level using standard keys. Next-generation sequencing was applied to soil eDNA to obtain earthworm sequence numbers and species diversity, and compared with numbers and diversity from hand sorting. We explore the diversity patterns generated by the two methods, assess the required sampling intensity, and examine sequence copy numbers, relative abundances and site occupancy proportions as potential proxy measures of abundance. Finally, we discuss the policy relevance of the findings and the potential effectiveness of earthworm eDNA for monitoring agricultural soil health and management interventions.

## 2. Methods

### 2.1 Study site

Earthworm hand-sorting and soil sampling for eDNA was performed at the University of Leeds commercial mixed farm in West Yorkshire, United Kingdom (53.870° −1.020°). Sampling was done at the end of March/early April 2018 when the weather conditions were broadly similar (overcast, sunny spells, 7–12°C). Three arable fields, conventionally cropped annually for more than 20 years, and a fourth field that had been arable for 6 years after 11 years in grassland, were sampled. Each field contained a pair of 72 m x 3 m strips of grass-clover ley that had been sown in 2015 as part of the SoilBioHedge project to examine the potential importance of hedgerow and field margin soils as biodiversity reservoirs for earthworms when converting to more regenerative agricultural management (Hallam et al., 2020). One of each pair of ley strips was separated from the field margin by a 13-m long and 90-cm deep stainless-steel mesh (0.104 mm pore size) curtain inserted vertically into the soil, extending approximately 10 cm above the soil, attached to a wooden frame (Figure 1). A 2-m wide strip of the field margin between the end of these ley strips and the hedges was kept fallow by regular glyphosate treatment and hand weeding (Figure 1). The other parallel ley strip in each field was continuous to the field margin adjoining the hedge, providing unrestricted access for soil biota from hedge-to-field. The ley strips were 33 months old at the time of sampling, and had been maintained by mowing and removal of biomass. The surrounding arable fields were annually ploughed and harrowed, and used to grow spring barley, winter barley and oil seed rape, respectively, during the three years of the growth of the leys (Guest et al., 2022). The arable areas of the fields received conventional chemical inputs (fertilisers, herbicides, fungicides, insecticides, molluscicides, and crop growth regulators), as detailed by Guest et al. (2022).

**Figure 1.**
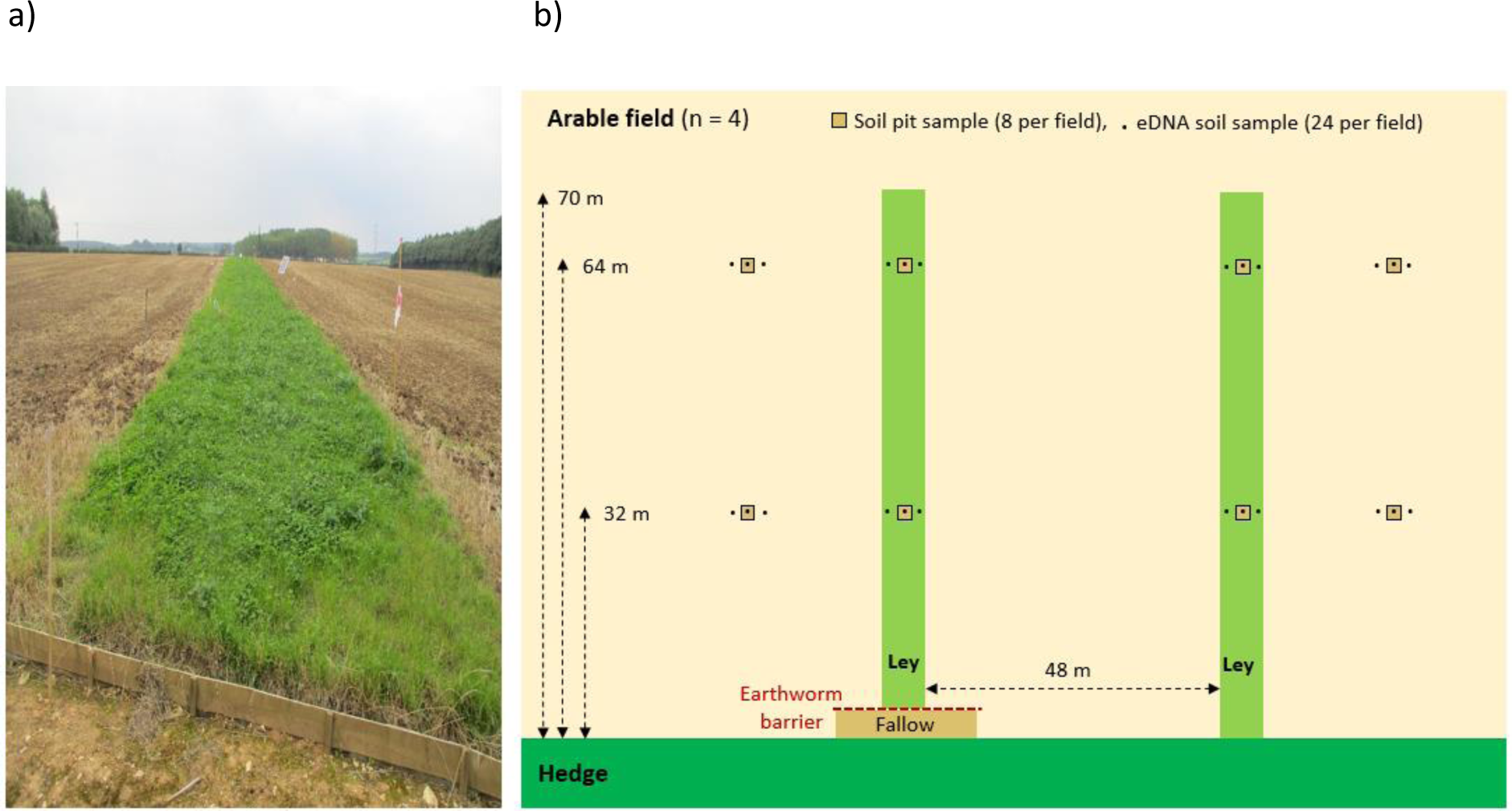
The experimental design and sampling protocol. (a) A ley strip with earthworm barrier in a barley field in autumn 2017 prior to sampling in April 2018, (b) the positions and numbers of samples taken per field, for both hand-sorting (*n* = 32 pits total) and eDNA (*n* = 96 soil samples total).

### 2.2. Soil sampling guided by findings from the SoilBioHedge experiment

Our sampling approach was guided by the earthworm population studies using soil pits and hand sorting, prior to the establishment of the ley strips in 2015 (Holden et al., 2019) and when the leys were one and two years old in April 2016 and 2017 (Prendergast-Miller et al., 2021). In the latter study, samples were taken at 2, 4, 8, 16, 32 and 64 m from the field edge, both along the paired ley strips and in parallel positions in the arable parts of the fields, to determine if earthworm species from the hedgerows and field margins that might be eliminated from the cropped areas colonised the ley strips. In SoilBioHedge over 260 soil pits were sampled, and 4,700 earthworms were identified to ecotype or species after preserving in ethanol by a professional earthworm taxonomist. This provides an exceptionally well-characterised study system in which to both evaluate earthworm eDNA methodology and establish the effects of increasing the duration of the leys from 2–3 years, which was outside the resources of the SoilBioHedge study.

Prendergast-Miller et al. (2021) found no evidence that the ley strips that contained the earthworm barriers were slower to recover earthworm numbers or species than those that were continuous to the field margin, nor was there any significant difference between earthworm abundance or biomass with distance from the field margin, both in the ley strips or arable field. As a result of these findings, in the present study, which lacked a large field team, soil sampling was only conducted at 32 and 64 m from the field edge to provide spatially representative replicate samples (Figure 1).

### 2.3. Soil environmental DNA sampling

In parallel to soil pit excavation (and prior to the handling of earthworms), soil blocks of approximately 5 x 5 cm and 15 cm deep were extracted for soil eDNA analysis using a pallet knife and a small trowel. Two were taken from 2 m either side of each soil pit and one was extracted from its centre, resulting in three soil samples for eDNA analysis for each earthworm-sampling soil pit (totalling 24 eDNA soil samples per field, Figure 1). The soil blocks were labelled in parallel with each soil pit, so that the eDNA and hand-sorting results were matched to the same field locations. Each eDNA soil block was transferred to a transparent, sealable plastic sample bag and carefully broken up within it so that any stones, large pieces of plant matter, earthworms or their cocoons could be removed. This process resulted in an average of ∼175 g of soil per eDNA sample. To prevent contamination from the tools used to take soil samples, these were cleaned and soaked in 10% bleach for at least 15 minutes before being reused. Disposable gloves were worn and changed between every sample extracted. At the end of each day, all equipment was cleaned again and hand tools were soaked in 10% bleach overnight. Excavated eDNA soil samples were stored at −20°C prior to eDNA extraction.

### 2.4. Earthworm sampling by soil pit excavation and hand-sorting

Pit digging for earthworm hand-sorting followed Holden et al. (2019) and Prendergast-Miller et al. (2021) in excavating an 18 x 18 x 15 cm deep block of soil. This was placed directly into a bucket with a sealable lid. After all earthworm pits and soil samples for eDNA analysis were taken from a field, the sealed buckets were transferred away from the field so that earthworms could be hand sorted and categorised into adults or juveniles. They were then preserved in 90% ethanol and kept refrigerated at 4°C prior to species identification. The numbers of preserved adults and juveniles per pit were counted and individual earthworm mass recorded following Prendergast-Miller et al. (2019; 2021). To ensure comparability between results of the present study and the previous work, we used the data archived by Prendergast-Miller et al. (2019) on earthworm numbers found from 0–15 cm depth, and as in the present study, included counts of damaged earthworms. Adults were identified to species under a dissecting microscope using morphological characteristics (Sherlock, 2018).

### 2.5. Earthworm environmental DNA extraction

Extractions of eDNA were performed using a modified protocol adapted from Taberlet et al. (2012, 2018). Soil samples were defrosted and thoroughly homogenised in their sealed bags by massaging the outside by hand. For each homogenised soil sample, 15 g subsamples were transferred into two 50 ml Falcon tubes, and 15 ml of freshly prepared saturated phosphate buffer (Na_2_HPO_4_, pH 8) was added using a bulb pipette to give a soil:buffer ratio of 1:1. For every three pairs of soil samples extracted, a blank extraction control, which consisted of phosphate buffer only, was included. The Falcon tubes were rotated for 15 minutes using a VWR tube rotator (VWR International Ltd, Lutterworth, UK), then centrifuged for 5 minutes at 3,130 X *g*, and 400 μl of the supernatant was pipetted off and processed using the Macherey–Nagel Nucleospin Soil kit (Macherey–Nagel, Düren, Germany) following the manufacturer’s instructions but skipping the lysis step. The resulting DNA extracts were diluted 1:5 with low TE buffer (10 mM Tris [pH8.0] 0.1mM EDTA) and stored in sealed tubes at −20°C prior to amplification. Extractions were all performed in a pre-PCR room inside laminar flow cabinets, which were thoroughly cleaned before and after use with 10% bleach and exposed overnight to UV-C light. Disposable gloves, weighing boats and utensils were changed between the handling of each sample and all equipment was wiped clean with 10% bleach and exposed to UV-C light overnight.

### 2.6. Environmental DNA amplification, purification and sequencing

*In-silico* analysis of two primer pairs proposed by Bienert et al. (2012), ewD/ewE and ewB/ewE (see Supplementary Materials 1.2 and 1.3) was performed to ensure suitability. The primers amplify short sequences (ewD/ewE ∼70 and ewB/ewE ∼120 base pairs respectively) of mitochondrial 16S rDNA (Bienert et al., 2012). The longer sequences amplified by ewB/ewE potentially reduces its capacity to detect more degraded eDNA.

For the initial amplification stage (PCR1), 2 µl of each diluted DNA extract was mixed with 1 µl each of the forward and reverse primers (5 µM), 10 µl of Qiagen Multiplex PCR Master Mix (Qiagen, Venlo, Netherlands) and 6 µl of molecular biology grade sterile water, totalling a reaction volume of 20 µl. All forward and reverse primers were tailed with Illumina sequencing primers. The PCR1 mixtures were prepared inside laminar flow cabinets in a pre-PCR room, and the reactions performed in individual 0.2 ml PCR tubes with sealable lids. In addition to extraction negatives, two PCR negatives containing ultrapure water instead of extracted DNA were included for every batch of reactions. For each batch, 20% of the extracted DNA samples were replicated (i.e. two PCR replicates produced for 20% of the samples) to enable reliability and sensitivity checks. After PCR1, the PCR product was visualised on a 1% agarose gel to ensure amplification had occurred, and to check for any contamination. PCR extracts were purified using AMPure XP beads (Beckman Coulter, Brea, CA). A second PCR step was then performed to add Illumina adapters and indexes, and product sizes were checked using a TapeStation (Agilent, Santa Clara, CA). The concentration of amplicons in each sample was then quantified using a fluorometer (BioTek Synergy LX Multimode Reader, Agilent, Santa Clara, CA) and samples were pooled together in groups of seven in equimolar amounts and a further bead clean undertaken. Each pool was quantified using qPCR and combined, resulting in one library for each primer pair. The BluePippin (Sage Science, Beverly, MA) system was used to size-select the final libraries to remove unwanted products, and the libraries were then combined into a single final library for sequencing on an Illumina MiSeq.

### 2.7. Bioinformatics and statistical analyses

Sequencing quality was assessed using FastQC and MultiQC (Andrews, 2010; Ewels et al., 2016). Sequences were trimmed using Trimmomatic (Bolger et al., 2014) to remove the Illumina adapter sequences from the data and trim lower-quality sequences. Reads were trimmed when the average Phred score dropped below 30 over a 4-base sliding window and reads below a minimum length threshold of 50 bp were discarded. MultiQC plots were then generated for the trimmed sequences to check that only high-quality data remained. The paired reads were aligned and converted to FASTA files using the FLASH alignment tool (Magoč and Salzberg, 2011), with maximum overlap and proportion of mismatches set to 150 bp and 0.1, respectively. The primer sequences were trimmed from the remaining sequences using the ‘trim.seqs’ command in the mothur software (Schloss et al., 2009), which was also used to label the sequences according to their respective primer pairs. After demultiplexing the sequences and producing separate FASTA files for each primer pair, USEARCH v9.2 (Edgar, 2010) was used to dereplicate the sequences, remove chimeric sequences and cluster highly similar sequences with 97% identity or greater. Unique mOTUs were generated for each primer pair and blasted against the NCBI nucleotide database, using a quality filter to only take forward hits with a maximum e-value of 0.00001 and 95% percentage identity (for the ewD/ewE primer pair this was increased to 97% percentage identity). The results were visualised using MEGAN 6 (Huson et al., 2016) and tables containing sequence ID and assigned taxon name were produced. Presence/absence and sequence number by sample matrices were then produced for each primer pair.

All statistical analyses were performed in RStudio using R version 4.0.1 (R Core Team, 2020) or Minitab. Analysis of the effects of the introduction of the ley strips, and their duration on juvenile, adult and total earthworm counts per square metre, combined data from Prendergast-Miller et al., (2019, 2021) and the present study. This was investigated using 2-way ANOVA tests with field treatment and duration of the study (years 1–3) and their interaction, as factors in the model, treating data from the different fields as independent replicates, and between means determined by Tukey tests. Effects of the third year of ley on the biomass of juvenile, adult, and total earthworms per m^2^ compared to under long-term arable field management was assessed using one-way ANOVAs.

Comparison of the percentage occupancy of each earthworm species in the 8 soil pits or paired triplicate soil samples per pit taken for eDNA analysis was calculated for each field. The average percentage occupancy across the four fields was compared by 2-way ANOVA with species and sampling method (hand counts versus ewD/ewE and ewB/ewE) and their interactions included in the analysis, using arcsine transformed data. Tukey tests were used to assess differences in percentage occupancy between species. Linear regression was used to investigate the relationship between hand-count occupancy data and that obtained using the best performing primer set (ewD/ewE).

To investigate the effect of the different sampling methodologies and agricultural treatments on earthworm species richness, a linear model was fitted using the function ‘lm’ with sampling method, ley versus arable and field ID as predictor variables, and species richness as the response variable. Model assumptions were checked and plotted using the ‘autoplot’ function in the ggplot2 graphics package (Wickham, 2011) and Tukey tests were carried out using the ‘TukeyHSD’ function. Bar charts were plotted using ggplot2 (Wickham, 2011).

To further examine possible community composition differences across the arable and ley treatments, and fields (and whether sampling methodology affected the results), non-metric multidimensional scaling (NMDS) was performed on the relative abundances from each sampling method using the ‘metaMDS’ function in the vegan package (Oksanen et al., 2013). The Bray–Curtis dissimilarity index was applied, and the ordinations were plotted with the environmental data overlaid using the ‘ordihull’ and ‘orditorp’ vegan functions. Shephard plots and stress-by-dimensions plots were also produced to check the suitability of the ordinations. One sample was removed from the ewB/ewE analysis as it skewed the NMDS2 axis and was deemed unrepresentative, probably because of low numbers of sequences. PERMANOVA tests were used to calculate the statistical significance of any potential differences in community dissimilarity brought about by the treatment groups, using the function ‘adonis2’ in vegan.

To evaluate the sampling intensities of each method and to check that the diversity present had been appropriately captured, species accumulation curves were plotted for each method, with one curve for each treatment type, using the ‘specaccum’ function in vegan. To investigate how closely the relative abundances of species in a sample compared across the different methodologies, scores per sample were plotted in stacked bar charts, overlaid with sequence numbers and abundances from traditional hand-sorting. Site occupancy proportions for each method were also calculated as potential proxies for overall species abundances.

## 3. Results

### 3.1. Abundance and biomass of adults and juveniles in arable and ley soils by hand sorting

In total, 718 earthworms were collected by hand-sorting, 594 of which were juveniles (82.8% of the total numbers) and 124 were adults (17.2%). Morphological identification of the adults enabled 119 earthworms to be assigned to species. Five earthworms could not be confidently identified due to damage during the sampling process. In total, eight species were identified, with *Allolobophora chlorotica* the most common (65 individuals), followed by *Aporrectodea rosea* (15) and *Lumbricus castaneus* (11) (Table 1). Of the identified adults, 88 were found in the 16 ley strip soil pits compared to 31 from the 16 arable field soil pits (goodness of fit χ^2^ = 26.4, 1 df, *p* < 10^-6^).

**Table 1.**
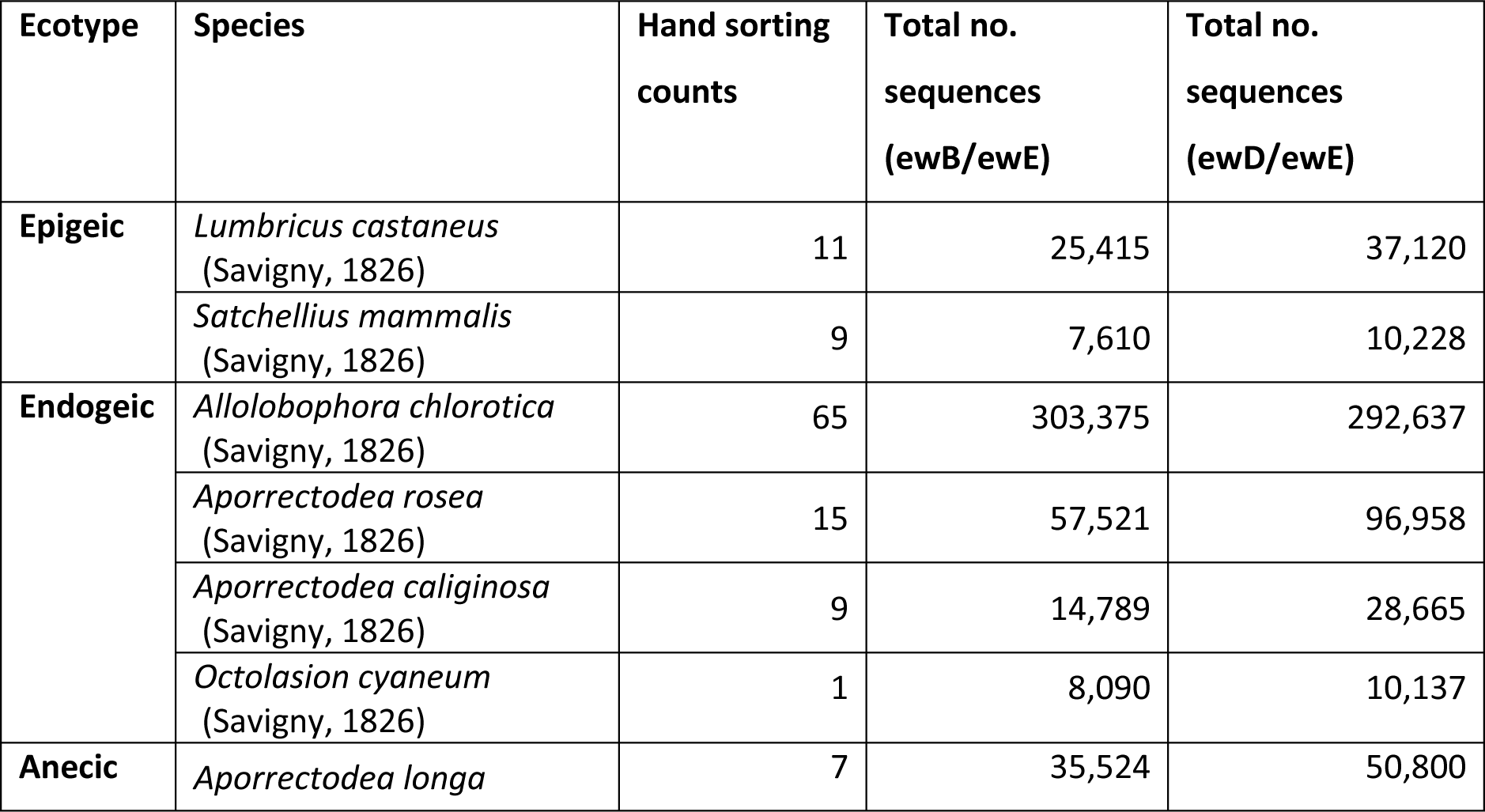
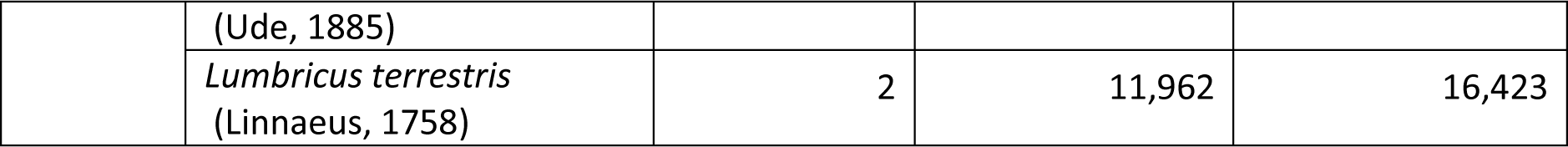
The total numbers of individuals and sequences found for each identified earthworm species by hand sorting and eDNA using the different methodologies.

There was no significant difference between the earthworm densities of the ley strips with and without the stainless-steel mesh barriers and fallow zone to the field margin (ANOVA: *F*= 0.18; 1,14 df, *p* > 0.05), nor was there a significant difference (ANOVA F= 1.19, 1,24 df, *p* > 0.05) between the two distances from the field margins (32 m vs 64 m). Data from the paired strips, and two distances, were pooled for subsequent analyses. This result was consistent with the suggestion of Prendergast-Miller et al. (2021) that regeneration of the earthworm populations in the leys was due to recruitment from the arable part of the fields, rather than by colonisation from the field margins. For the leys there was no significant difference between the four fields in earthworm densities in 2018 (Supplementary Material, Figure S1). However, in the arable areas, one field (Hillside) that most recently had been in grassland (2009) had significantly higher total earthworm densities (mean 300 m^-2^) than the arable field (BSSW) with the lowest (mean 46 m^-2^) earthworm density (natural log transformed data, ANOVA: *F* = 4.19, 3,12 df, *p* < 0.05).

The density of adult and juvenile earthworms per square metre in arable fields and ley strips in 2018 (Figure 2a) was compared with the data published for the previous two years of the field experiment (Prendergast-Miller et al., 2019; 2021). These values were visually benchmarked against the average total earthworm density of the field grassy margins beside hedges and four adjacent fields under permanent grassland for silage or sheep grazing that had been quantified from 2015–17 and had showed no significant difference between years (Holden et al., 2019). Importantly, our April 2018 data for the 3-year ley showed surprisingly large and significant increases (Tukey test *p* < 0.05) in both juvenile and total earthworm densities compared to the two-year leys which had densities similar to the grassy field margins and adjacent permanent grasslands (Figure 2a).

**Figure 2.**
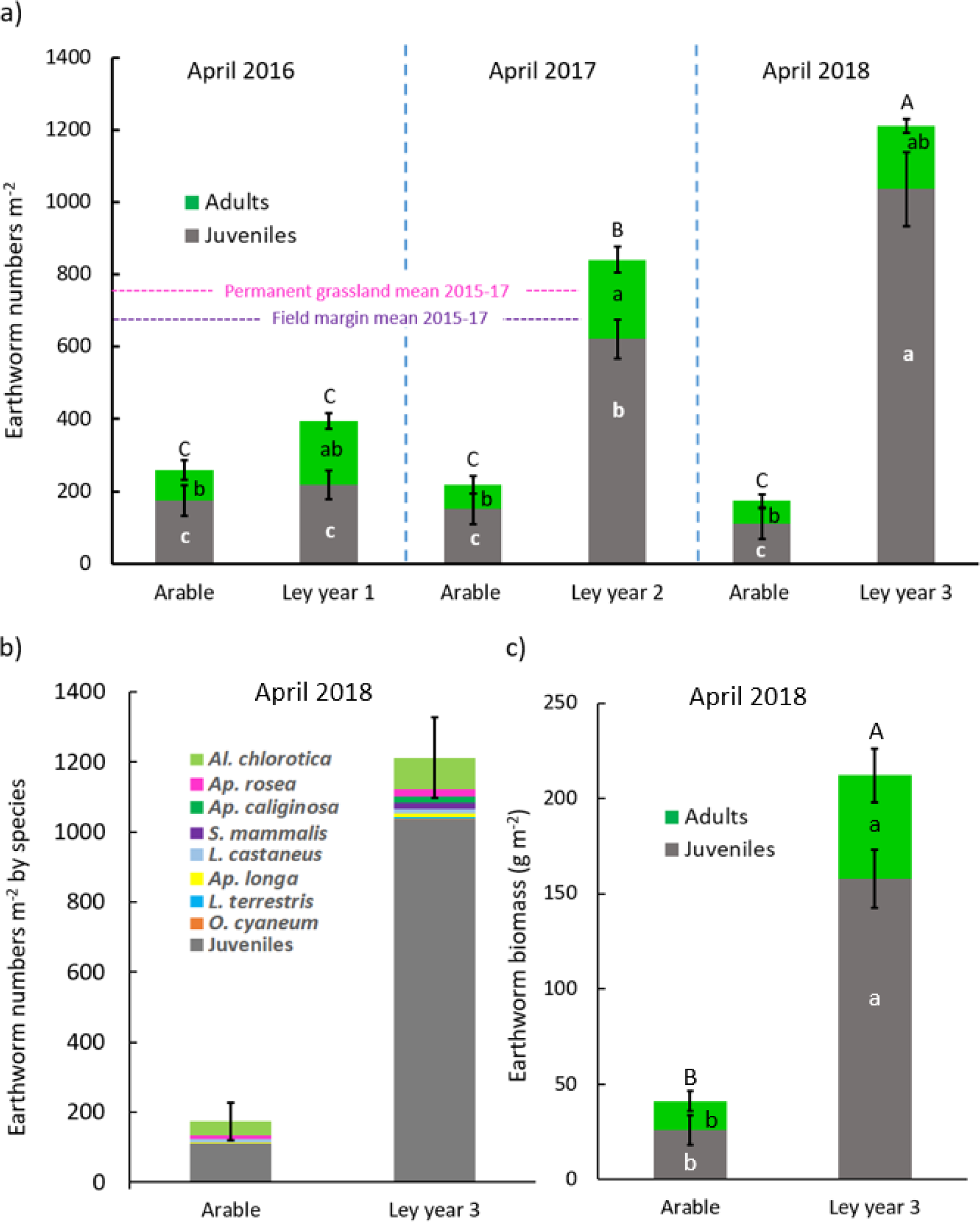
(**a**) The number of earthworms as juveniles and adults per square metre in the arable fields and ley strips over 3 successive years. April 2016–April 2017 data are analysed from Prendergast-Miller et al. (2019), and for April 2018 are from the soil pit results of the present study (with standard errors of the means for *n* = 4 fields). (**b**) The mean numbers of earthworms in April 2018 identified to species, with standard error of the mean total numbers of earthworms shown (*n* = 4 fields). (**c**) The mean biomass of juvenile and adult earthworms per m^2^ in 2018, with standard errors of means (*n* = 4 fields). Results of Tukey tests within each panel **a**–**c** are shown by letter codes, with stacked bars sharing the same uppercase letters above them being not significantly different (Tukey test *p* <0.05), bars sharing the same lowercase black letters show no significant differences for adult earthworms, and bars sharing the same lowercase white letters show no significant difference for juveniles (*p*> 0.05).

Total earthworm numbers per m^2^ were significantly regenerated by leys compared to the arable controls (ANOVA: *F* = 28.9 1,23 df, *p* <0.001). There was a significant effect of year (*F* = 12.41 2,23 df, *p* <0.001) which was driven by the interaction of the increasing regenerative effect of the leys over time (*F* = 18.8 2,23 df, *p* <0.001). In each successive year of ley after 2016, both total earthworm numbers and juveniles increased significantly (Tukey test *p*<0.05), whereas in the arable parts of the fields there was no difference between years (*p*> 0.05). By contrast, with adults there was no significant difference in density between years (*p* >0.05), no interaction between ley and year of study (*p* >0.05). Although there was an overall significant increase in adults in the ley compared to arable parts of the fields (ANOVA: *F* = 33.4 1, 23 df; *p* < 0.001), this difference was only significant in 2017 (*p* <0.05) and not 2018. The total earthworm abundance in the 3-year leys was, on average, nearly 7 times higher than in the arable parts of the fields, reaching over 1,200 individuals per m^2^ (Figure 2a). There were over 9 times more juvenile earthworms per square metre in the 3-year ley than the arable sites, confirming the suitability of the ley habitat for earthworm reproduction.

At the species level (Figure 2b), adult earthworms in April 2018 were dominated by *Al. chlorotica* (42 and 94 individuals m^-2^, in the arable and 3-year leys respectively), and in the leys the next four most abundant species (*Ap. rosea, Ap. caliginosa, S. mammalis,* and *L. castaneus*) had similar abundances (15–20 individuals m^-2^). The least abundant species in the 3-year leys were *Ap. longa,* (10 individuals m^2^), *L. terrestris* and *O. cyaneum* (2–4 individuals m^-2^). Only half the species found in the 3-year leys were recorded in the arable sites, with *Ap. caliginosa, S. mammalis, L. terrestris* and *O. cyaneum* not being detected.

Earthworm biomass showed a five-fold increase in the 3 year leys compared to the arable parts of the field (Figure 2c), with total, adult and juvenile earthworm biomass all being significantly increased (ANOVA: total biomass *F* = 91.4 1,6 df, *p* < 0.001; adults *F* = 9.9 1,6 df, *p* < 0.05; juveniles *F* = 67.5 1,6 df, *p* < 0.001). Because of their larger size, adults in the 3-year leys contributed a greater proportion to total biomass (25%, Figure 2c) than to total numbers (14%, Figure 2a).

#### Environmental DNA sequences

MiSeq sequencing yielded a total of 9,464,039 paired reads from the eDNA samples (including sequences from both primer pairs but not negatives and repeats). After the initial trimming based on sequence length and quality, 4,693,446 paired reads remained. On average, 24.9% of reads were removed during quality control. The sequences then underwent further quality filtering and selection, including the removal of chimeras and applying stringent BLAST criteria, which reduced the sequence numbers further.

After the bioinformatics clean-up and filtering steps had been completed, the total number of sequences classified to species level was 794,350 for ewB/ewE and 707,750 for ewD/ewE (not including the sequences obtained from negatives and repeats). For ewB/ewE, 464,287 were earthworm sequences and 330,063 were from the closely related *Enchytraeidae*, compared with 542,974 earthworm and 164,776 enchytraeid sequences for ewD/ewE. The eDNA primer pairs each detected the same eight earthworm species as found with hand sorting, with ewB/ewE identifying an additional nine enchytraeid species compared with eight for ewD/ewE. Of the eight earthworm species identified by both methods (Table 1), the species with the highest sequence numbers overall was *Al. chlorotica* (with 303,375 and 292,637 sequences found by ewB/ewE and ewD/ewE, respectively), followed by *Ap. rosea* (57,521 and 96,958) and *Ap. longa* (35,524 and 50,800).

#### Comparison of percentage site occupancy by species by hand sorting and eDNA

The percentages of soil pit samples in which each of the 8 species were recorded by hand sorting or by 3 replicate soil samples from the match location using eDNA methods were compared (Figure 3). Although both eDNA primer sets tended to show higher detection rates than the hand sorting method,

**Figure 3.**
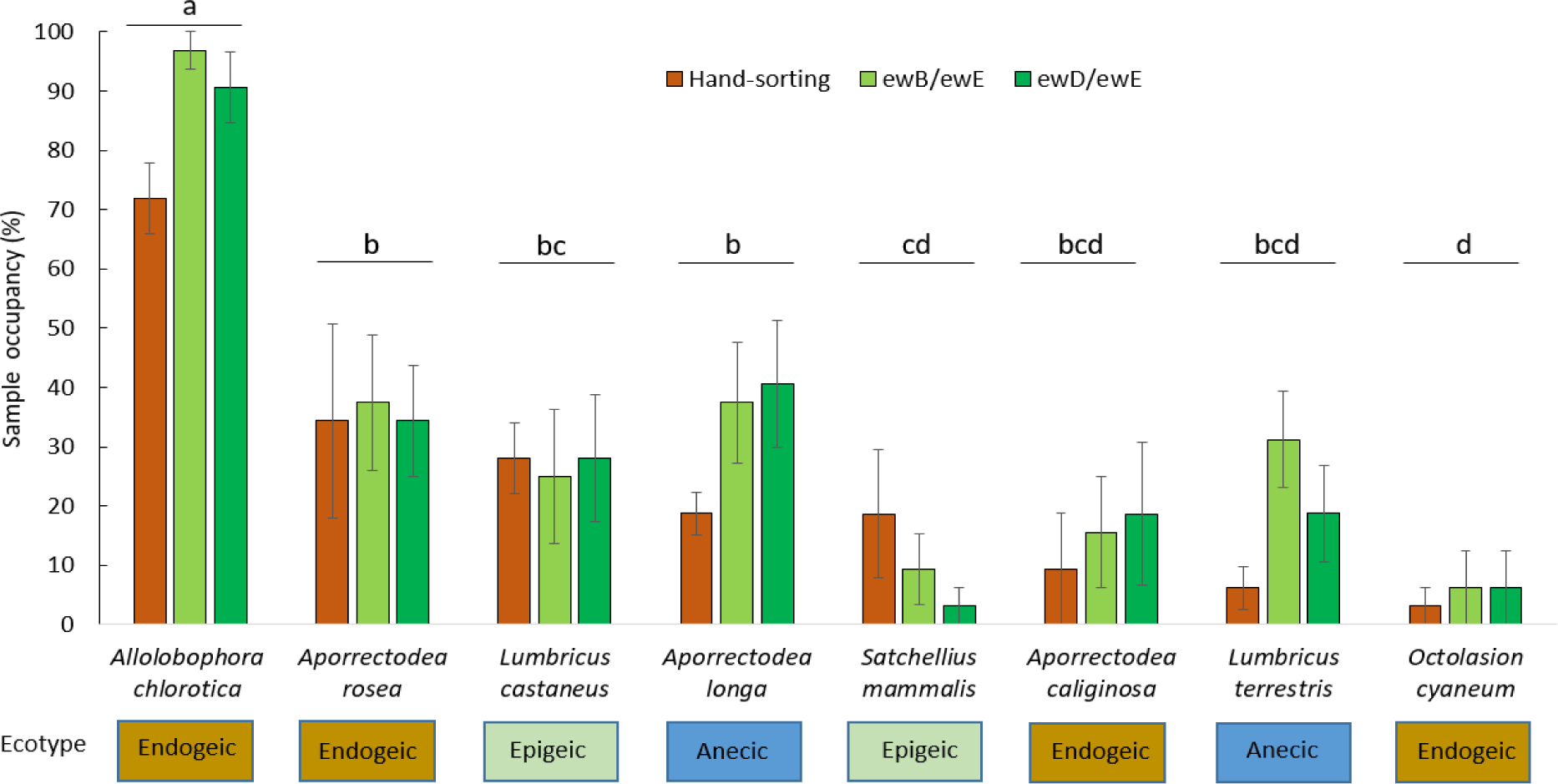
Mean percentage of occupancy of the 8 earthworm species found by hand sorting and identification of adults, and by eDNA using ewB/ewE and ewD/ewE. The values are means of the average occupancy of 8 soil pits per field, or triplicate eDNA soil samples matched to each pit, pooling arable and ley samples (*n* = 4 fields in each case), with standard errors of the mean shown. The ecotypes are indicated for each species. Species sharing the same letter codes are not significantly different (*p* > 0.05) in Tukey tests performed on arcsine transformed data averaging across the three methods of detection.

ANOVA analysis on arcsine transformed percent occupancy data found no significant difference between hand sorting and the two eDNA methods (*p* > 0.05), nor was there a species by method interaction (*p* >0.05). However, as would be expected, there were significant differences (Figure 3) in the occupancy detected for the different species (*F* = 21.7 7,72 df; *p* < 0.001).

For the primer pair ewD/eWE, which detected the most earthworm eDNA sequences, and tended to provide the highest site occupancy rates in Figure 3, we conducted linear regression between the percentage of soil samples containing eDNA sequences for each earthworm species and the percentage of soil pits in which each species was found (Figure 4). The best fit line was *y* = 1.1684*x* + 4.588, *R*² = 0.81 and *p* < 0.01. The endogeic ecotype species most closely fitted the regression line, with the deep burrowing anecic ecotypes, especially *L. terrestris,* lying above the fitted line, and the epigeic, surface dwelling species, lying below it.

**Figure 4.**
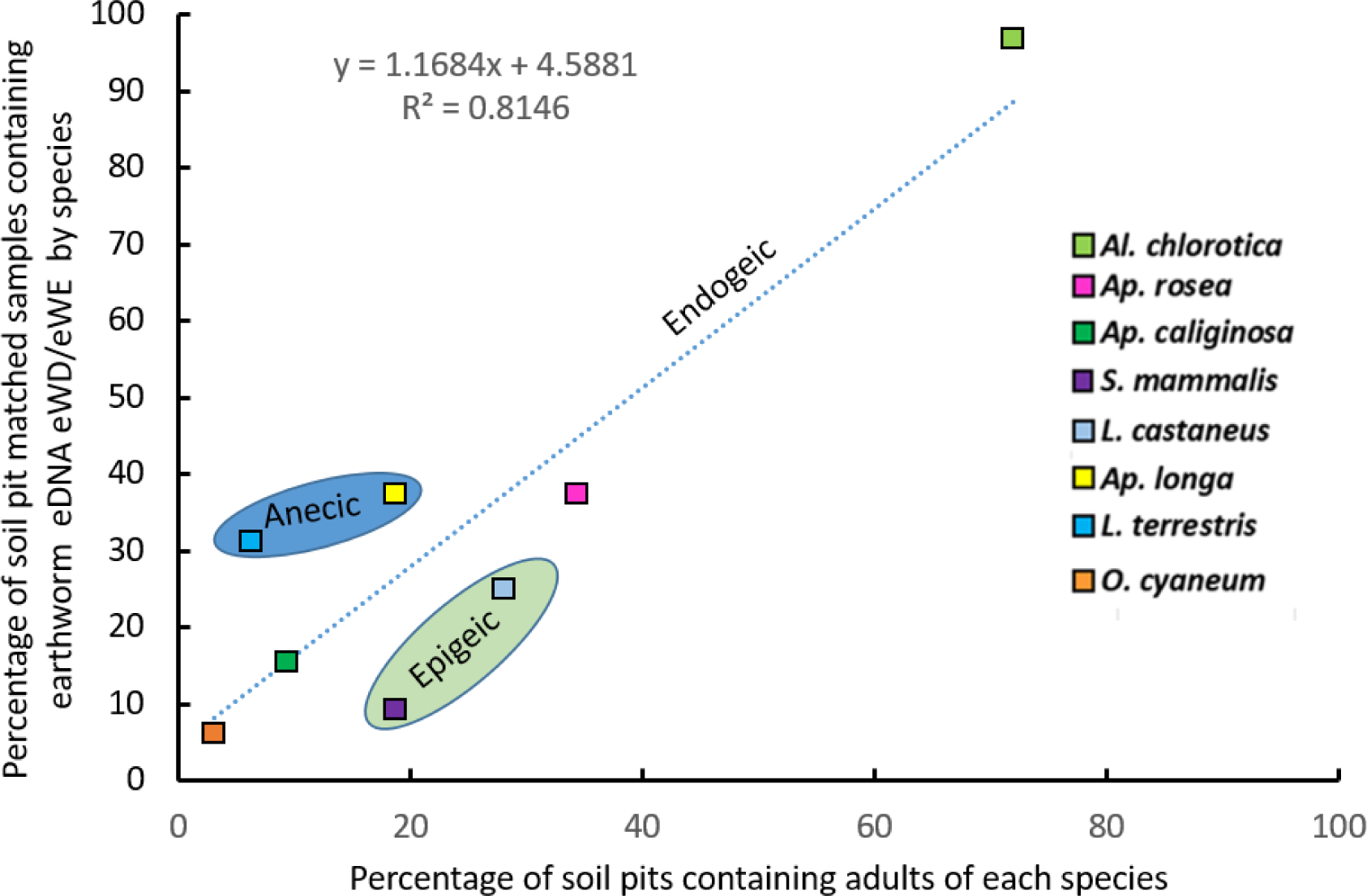
Linear regression between the percentage of soil pits containing adults of each of the 8 earthworm species, and percentage of soil samples containing eDNA sequences of these species using ewD/ewE and pooling data from all sampling sites in April 2018. The three ecotype groups of earthworms are overlain as differences in their sizes, behaviour and soil occupancy may affect their detection by hand sorting and eDNA. Full species names are given in Table 1.

#### Comparison of eDNA sequence numbers by species, and species abundances by hand sorting

For the primer pair ewD/ewE, the percentage of total sequences by species was plotted against the percentage abundance of the identified adult earthworms of each species (Supplementary Materials, Figure S2), pooling the data across all fields and arable and ley treatments as in Table 1. This showed that there was a highly significant correlation (*R*^2^ = 0.96; *p* <0.001) between the relative abundance of eDNA sequences for each species and the percentage of the adult earthworms that each species contributed to hand-sorting records. Furthermore, the slope of the line indicated an almost 1:1 ratio between the two measures. A similar, highly significant relationship (*R*^2^ = 0.98; *p* < 0.001) was also found for ewB/ewE sequences (Supplementary Materials, Figure S3). The similarity in adult earthworm species abundances m^-2^ and the abundance of eDNA sequences using ewD/ewE for each species averaged for arable and ley samples are shown in Supplementary Materials, Figure S4, and the data of the individual soil samples shown in Figure S5.

#### Sampling method comparison, and management effects (arable vs ley) on earthworm species diversity

Although both eDNA primer sets and the hand-sorting method identified the same eight earthworm species, the mean earthworm species richness per pit sample was significantly affected by the sampling method (ANOVA: *F* = 12.79, *df* = 2, 87, *p* < 0.001). Hand-sorting gave lower earthworm species richness than both the ewB/ewE (Tukey multiple comparison test, *p* = 0.006) and ewD/ewE (*p* < 0.001) eDNA methods (Figure 5). There was no significant difference in mean earthworm species richness between the two eDNA methods (*p* = 0.17).

**Figure 5.**
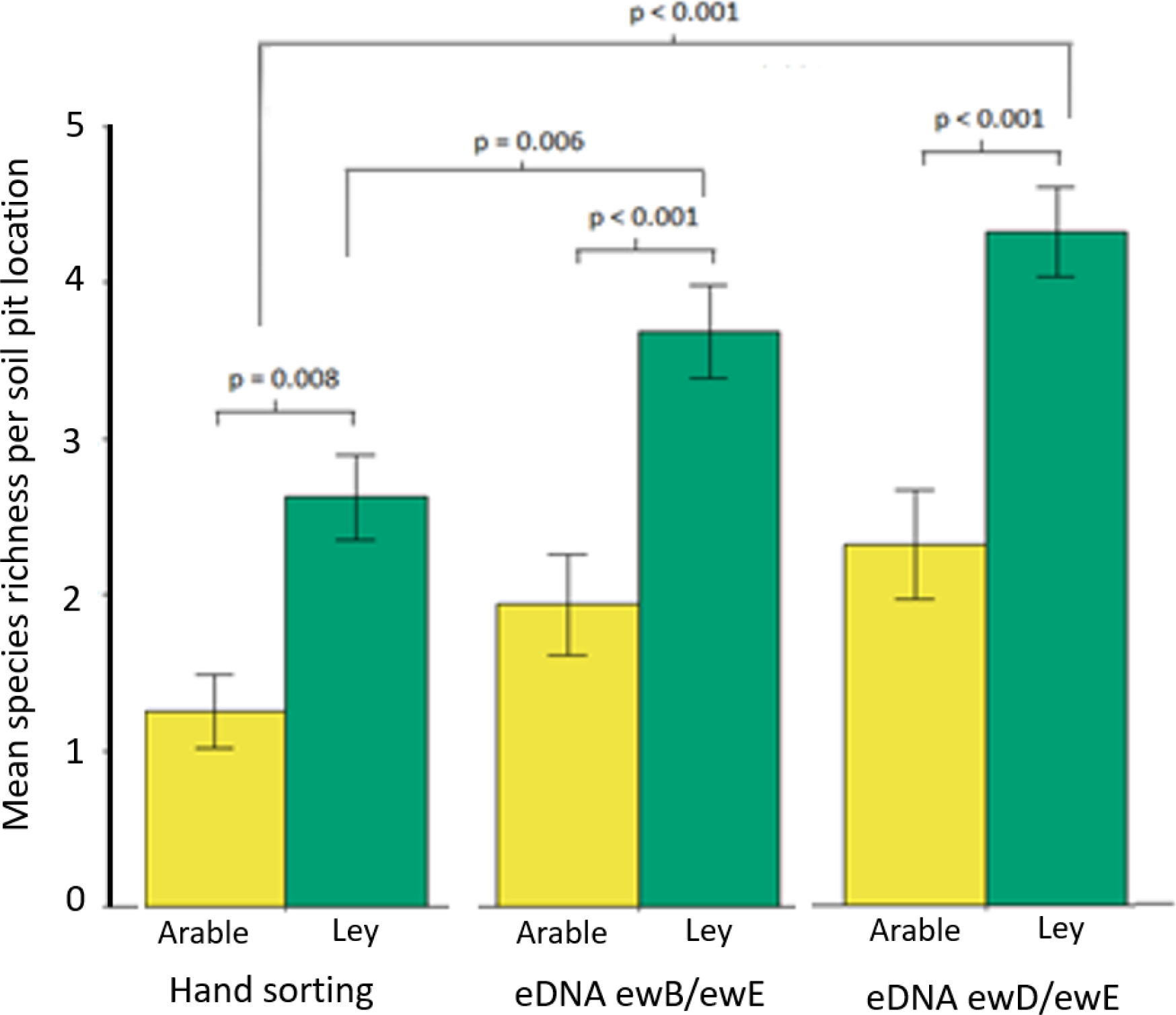
Mean earthworm species richness per pit (± standard error) determined using hand sorting (*n* = 16) compared with eDNA primer pairs ewB/ewE or ewD/ewE), for arable (*n* = 48) and ley (*n* = 48) parts of the four fields. Significant differences between the methods and between arable and ley for each method are shown by the *p* values.

The three-year ley approximately doubled species richness (Figure 5) compared to the arable parts of the fields, giving an overall highly significant effect averaging across results of the three sampling methods (ANOVA: *F* = 57.78, df = 1, 87, *p* < 0.001). Tukey-test comparisons found that the increase in species richness in the leys detected by hand sorting was slightly less significant (*p* = 0.008) than that detected by both eDNA methods (*p* < 0.001). Location-specific effects were also observed; earthworm species richness differed among the four fields (ANOVA: *F* = 5.65, df = 3, 87, *p* = 0.001), which was driven by significantly higher mean species richness in one field that had most recently been in grassland for 11 years and then had been cultivated for the previous 9 years compared to the long-term (>20 years) arable field with the lowest earthworm numbers (Tukey multiple comparison test, *p* < 0.001). The three long-term arable fields showed no significant differences in earthworm species richness.

#### Species accumulation curves with sample numbers

For both hand sorting and the two eDNA methods, the species accumulation curves (Figure 6) indicated that the numbers of soil pits and eDNA samples taken were sufficient to detect all the common species in the ley and arable areas of the four sampled fields. In all cases, the ley samples showed steeper initial rates of cumulative species gain per soil sample than the arable samples. For hand-sorting, the arable treatment curve reached a plateau at four species, from the 16 soil pits (soil volume of approximately 72 litres). Similarly, the arable curve levelled out at six species for the ewB/ewE eDNA method as *O. cyaneum* and *S. mammalis* were not detected in the 48 samples analysed (approximately 18 litres of soil). In contrast, all eight species of earthworm were recorded in the arable control treatments using the ewD/ewE method, although it took more samples for the arable species accumulation curve to plateau compared with the ley treatment curve.

**Figure 6.**
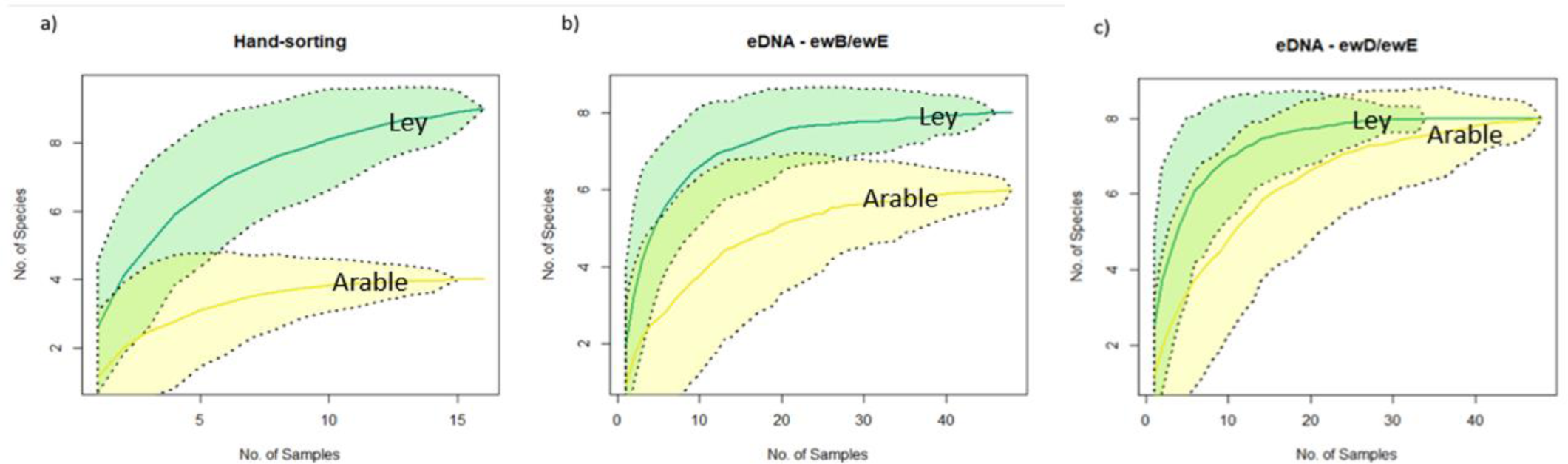
Species accumulation curves (a) by numbers of soil pits (each ∼4860 cm^3^) and hand-sorting, or soil sampling (375 cm^3^) for eDNA analysis by (b) ewB/ewE primers or (c) ewD/ewE primers respectively. Green lines represent curves for the ley treatment and yellow lines are for the arable treatment. Coloured areas around the lines with the dotted line borders indicate 95% confidence intervals.

Importantly, the ewD/ewE eDNA method detected earthworm species in the arable parts of the field which were not detected by hand sorting, but which must have provided the diversity that was recruited into leys, giving their more diverse communities. However, to detect these low abundance species in the arable soils appears to require over 40 samples for eDNA analysis (approximately 15 litres of soil, from which 600 g total was extracted after homogenization of each sample). The increased sensitivity of the eDNA approaches is notable when looking at the overall soil volumes analysed in comparison with hand sorting. For hand sorting a total volume of 0.156 m^3^ of soil was sorted across 32 soil pits, in comparison to a total volume of 0.036 m^3^ analyses across 96 eDNA samples. The fact that eDNA approaches detected more species in less soil (only 23% of the soil pit volume sampled in hand sorting) indicates its greater sensitivity.

#### Multivariate community analysis of earthworm populations detected by hand sorting and eDNA sequences

Using Bray–Curtis dissimilarity calculations, NMDS ordinations successfully reached convergent solutions after 40 iterations for all three methods. With two-dimensional scaling, the reported stress levels were 0.130 for hand-sorting, 0.156 for ewB/ewE and 0.154 for ewD/ewE. Visual inspection of the NMDS ordinations suggest considerable overlap between communities belonging to both the arable and ley treatments (Figure 7), indicating that communities in the ley treatment were not clearly dissimilar from arable communities at this stage. This was supported by subsequent PERMANOVA tests, which did not find significant differentiation between the earthworm communities according to field management treatment (for hand-sorting pseudo-*F* = 1.37, *p* = 0.25; ewB/ewE pseudo-*F* = 1.33, *p* = 0.23; ewD/ewE pseudo-*F* = 2.21, *p* = 0.07). The field management treatment groups were also found to have homogeneous dispersions that did not differ significantly (betadisper tests, for hand-sorting *p* = 0.59, ewB/ewE *p* = 0.29, ewD/ewE *p* = 0.33). There was some distinction between communities between the four fields observed in the hand-sorting method (PERMANOVA; pseudo-*F*= 2.35, *p* = 0.02), although this was not observed by the two eDNA methods (ewB/ewE pseudo-*F* = 1.32, *p* = 0.21; ewD/ewE pseudo-*F* = 1.32, *p* = 0.19; see Supplementary Materials 2.2 Figures S6-S8 for NMDS plots overlaid with field name groups).

**Figure 7.**
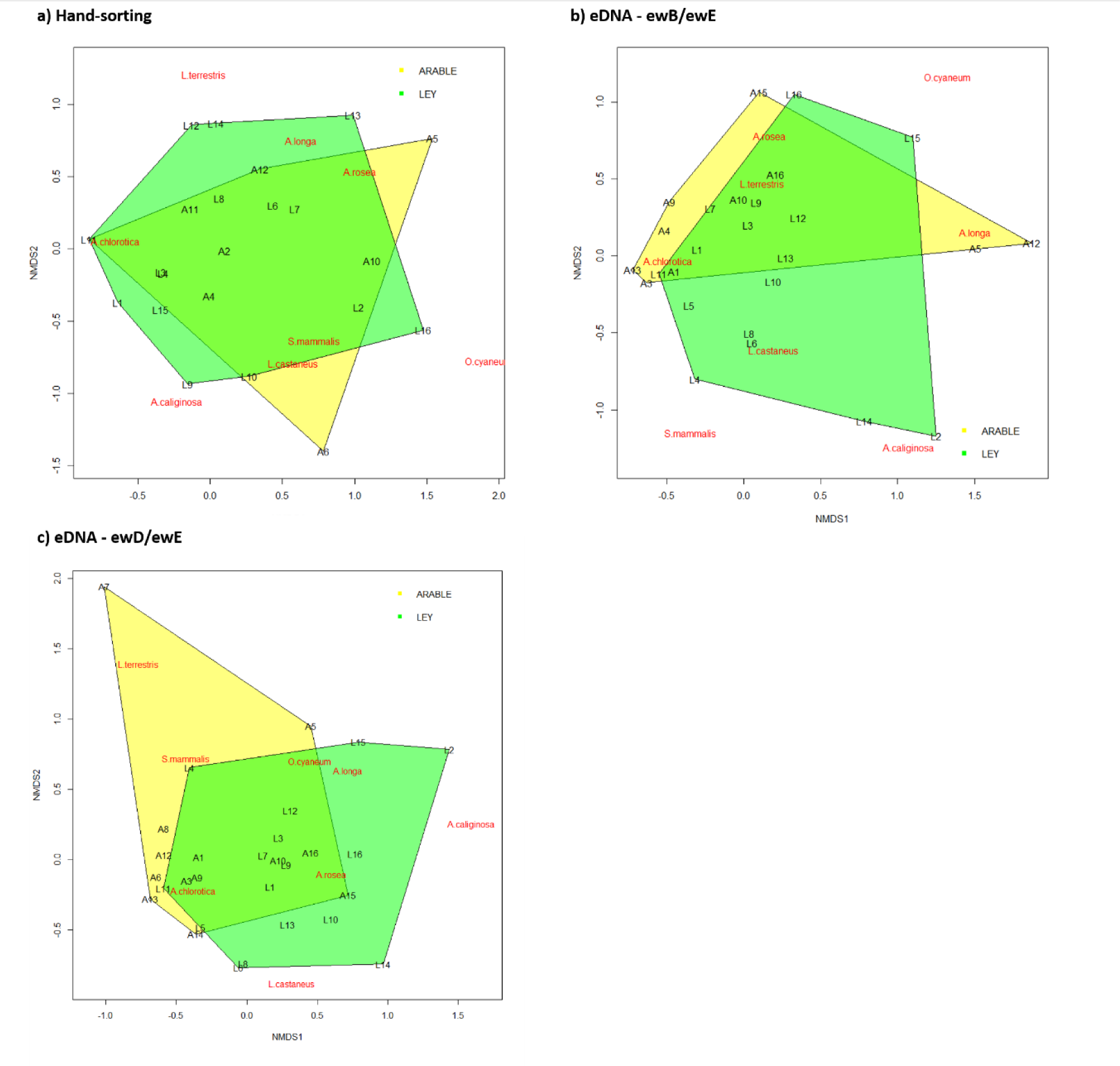
NMDS ordinations for (a) hand-sorting, (b) eDNA sampling using ewB/ewE primers and (c) eDNA sampling using ewD/ewE primers. The site scores in ordination space are represented by the site labels (arable sites = ‘A1’, ‘A2’ etc., ley sites = ‘L1’, ‘L2’ etc.), and the species labels are positioned at the weighted average of the site scores. The polygons connect the vertices of points made by the communities in the ley (green polygons) and arable (yellow) treatments.

## Discussion

Our study establishes rapid regeneration of earthworm populations in three-year grass-clover leys introduced into conventionally managed, arable fields that had been intensively cultivated and cropped for decades, and demonstrates, for the first time, that eDNA sampling is effective for monitoring earthworm species and appears to provide an indication of their relative abundances in arable fields and leys. For measuring earthworm species richness, both eDNA primer pairs performed well compared with hand-sorting, identifying the same 8 species, but almost consistently detecting more species per sample, despite using much less soil than standard soil pits. They revealed the same pattern of increased species richness in the 3-year leys compared to permanent arable parts of the fields, and the relative abundance of species assessed by eDNA sequence numbers showed strong linear correlations with hand-sorted counts of adults of the same species. The species accumulation curves indicated that the pit sampling replication for both traditional hand-sorting and eDNA soil sampling was appropriate for this study system, and effectively characterised the earthworm biodiversity in both arable and ley samples. This is corroborated by the SoilBioHedge research on the same fields in the three years preceding the present study. In that research, a team of more than 12 people surveyed a total of 36 soil pits in the field hedges, 36 in grassy field margins inside hedges, 72 in the arable fields, and 96 pits in the 1–2-year-old ley strips, together with 24 pits in four adjacent permanent pastures, totalling over 260 soil pits (Prendergast-Miller et al., 2021). A total of 1,329 adult earthworms were identified by an expert, with the 8 species found in the present study comprising 99.1% of the identified individuals, three other species occurring as adults with just one individual (*Eisenia fetida*), two individuals (*Dendrodrilus rubidus*) and nine individuals (*Murchieona muldali*) out of 4,700 adults and juveniles counted (Prendergast-Miller et al., 2021). Given the rarity of these other species, which may have occurred outside the arable and ley areas, and the intensity of sampling required to detect them, the eDNA methodology appears to be highly effective for earthworm biodiversity monitoring.

Both eDNA primer pairs gave consistent results and captured similar pictures of earthworm diversity, so both are potentially useful in wider-scale monitoring programmes. As predicted by Bienert et al. (2012), ewB/ewE provided better species resolution but amplified a greater proportion of *Enchytraeidae* sequences. Although we chose to focus solely on earthworms in this study, enchytraeids are also functionally important biota and might serve as indicators of soil health (Koutika et al., 2001; Marinissen and Didden, 1997; Pelosi and Römbke, 2016), so future surveyors employing eDNA sampling of soils for earthworm biodiversity may wish to utilise the ewB/ewE primers in order to include these taxa. However, enchytraeids may be less well represented in the sequence databases than earthworms, so may require additional work to establish a suitable reference database. For surveys looking solely at earthworms the ewD/ewE primers may be preferable, as these amplified more earthworm sequences overall.

The identification of more earthworm species per sample by the eDNA methods than by hand sorting may in part be due to detecting juveniles of species, which cannot be reliably identified morphologically. Over 80% of earthworms collected through hand sorting were juveniles, which is a large potential reservoir of diversity that is excluded from the hand-sorting results. The lower species diversity obtained through traditional hand sorting may also be due to worms fleeing the pit areas as the soil blocks are excavated. Earthworms are sensitive to vibration and the larger vertical burrowing anecic worms, whose burrows can extend well below the depth of the excavated pits, may retreat to evade capture (Pelosi et al., 2009; Singh et al., 2016). The eDNA methods were markedly better at detecting the two larger anecic species *A. longa* and *L. terrestris* (for *A. longa*, site occupancy measured by hand-sorting was 18.8% compared with 37.5% and 40.6% for the eDNA methods; for *L. terrestris*, hand-sorting site occupancy was just 6.3% compared with 31.3% and 18.8% for eDNA). These two species have different feeding and burrowing behaviours, with *L. terrestris* forming permanent vertical burrows that would be expected to give an eDNA “hot spot” if included in a soil sample, and actively forage for fresh plant litter on the surface (Hoeffner et al, 2022), potentially leading to the low count number but high frequency of detection by eDNA. Our finding of one eDNA sample with over 11,000 copies of *L. terrestris* eDNA and of large numbers of samples having only very low counts is consistent with this anticipated behavioural and soil occupancy effect. By contrast, *A. longa* behaves more like an endogeic species, consuming decayed plant material and producing a more extensive burrow system (Hoeffner et al., 2022), and in the relationship between percentage soil occupancy assessed by eDNA and by hand sorting (Figure 4), it was positioned close to the best-fit line to which the endogeic species fitted as a group.

To increase the recovery of deep-burrowing anecic species, researchers have added chemical expellant to soil pits, such as allyl isothiocyanate or formaldehyde (for example, Crittenden et al., 2015; Holden et al., 2019 and Prendergast-Miller et al., 2021). Due to the need to rapidly process samples for eDNA, we did not include this step, which requires observing the pits for 30 minutes. The chemical extraction step has also not been used in farmer-participatory sampling such as the #60minworms and #30minworms studies (Stroud, 2019; Stroud and Goulding, 2022). In the SioBioHedge study, allyl isothiocyanate was added to soil pits and both *A. longa* and *L. terrestris* were found at densities of 3.6 m^-2^ in the arable parts of the fields and 5.4 m^-2^ in the leys, averaging across 2016–2017 (Prendergast-Miller et al., 2021). In our study, not using the chemical, we found *A. longa* in the arable and ley parts of the same fields in 2018 at densities of 4.2 m^-2^ and 10.5 m^-2^, respectively, but found no *L. terrestris* in the arable pit samples, but at densities of 4.2 m^-2^ in the ley, so not using the chemical does not appear to have substantively changed the results.

The only species that was found to have equal or slightly lower site occupancy percentages when measured using eDNA compared with hand-sorting was the epigeic worm *S. mammalis* (6.3% and 18.8% for the eDNA primer pairs compared with 18.8% for hand-sorting). Given that all methodologies found *S. mammalis* to be relatively rare at the study site, this may simply be due to the heterogeneous spatial distribution of earthworm populations in soils (Valckx et al., 2011). However, Bienert et al. (2012) also reported underrepresentation of epigeic species by eDNA sampling. They attributed this to the limited number of samples taken and not sampling the top few centimetres of soil where epigeic earthworms would be most active. However, although our study avoided these limitations, we still observed the same outcome, which may indicate other possible causes. It has been shown previously that numerous factors can affect eDNA stabilisation and degradation rates in soils, including moisture levels, pH, temperature, microbial activity, soil type and chemistry (Barnes and Turner, 2016; Harrison et al., 2019; Pietramellara et al., 2009; Sirois and Buckley, 2019). Exposure to UV light at the soil surface might be expected to accelerate eDNA degradation, although evidence for this appears to be mixed (Harrison et. al., 2019). Many environmental variables vary with soil depth, and it is likely that the surface soil may experience conditions that increase eDNA degradation rates so that surface-dwelling species are less represented. Furthermore, it was only the longer ewB/ewE primer pair that showed lower site occupancy, and we would expect longer DNA fragments to be more susceptible to degradation. Future studies may shed light on whether the lower detection of epigeic species here is consistent, and a result of eDNA breakdown at faster rates near the soil surface. More research is needed on how eDNA stabilisation and degradation rates may be affected by soil depth, particularly given the potential role of earthworm-mediated bioturbation in the transport of DNA molecules throughout the soil profile, as demonstrated by Prosser and Hedgpeth (2018), and the effects of earthworm casting and litter-feeding on the surface (Hoeffner et al., 2022).

The capacity for eDNA to persist in the environment has led to ongoing debate over the extent to which eDNA sampling is measuring current or past populations (Thomsen and Willerslev, 2015; Sirois and Buckley, 2019). Our study shows clear differences in soil eDNA profiles after three years under different management treatments and these patterns are similar to those revealed by traditional sampling techniques. This shows that eDNA sampling is sensitive enough to pick up fine-scale differences in earthworm communities in a relatively short time period, despite any ‘background’ eDNA that may be present. It could therefore be a useful tool for gauging the success of new soil management regimes that have been implemented to improve on-farm soil biodiversity.

We cannot, however, definitively rule out that some of the sequences in our totals may originate from background eDNA deposited by earthworms that were no longer present. It is possible, for example, that for the ewD/ewE primer pair the small numbers of sequences of *S. mammalis* and *O. cyaneum* in the arable samples may be due to eDNA from populations recorded in the previous years (Prendergast-Miller et al., 2021), as these species were not detected in 2018 either by hand-sorting or the longer ewB/ewE primer pair (which would not have detected shorter, more degraded eDNA fragments). However, it is also plausible that the eDNA detected the presence of these relatively rare earthworms as juveniles, as they comprised only 4% and 0.4% of the adult earthworm populations recorded across all the fields and sites sampled by Prendergast-Miller et al. (2021). Future research is needed to better understand the degradation rate of eDNA in soils, and ways to account for background effects (Thomsen and Willerslev, 2015). A recent study by Marshall et al. (2021) offers a useful starting point for this, describing a method for estimating the age of eDNA based on accompanying eRNA sampling and analysis of the eDNA:eRNA ratio. Utilising techniques like this may prove useful in agricultural soil eDNA surveys, as they would allow the surveyor to account for variation in eDNA degradation rates that might be brought about by different agricultural practices (Sirois and Buckley, 2019; Foucher et al., 2020).

Sampling intensity and effort is another important aspect to consider if earthworm eDNA sampling is to be more widely used in soil monitoring schemes (Dickie et al., 2018). The species accumulation curves indicated that an appropriate sampling replication was carried out for this study system but, for accurate sampling of larger sites and whole farms, more samples would be needed. The sampling intensity required for a complete inventory of all species is clearly much greater than that required to detect the widespread and dominant species. From a soil health and functional perspective, it seems unlikely that rare earthworms deliver substantial benefits other than as contributors to biodiversity per se.

Further work to develop a standardised eDNA sampling scheme would be invaluable. This should be done across a range of agricultural sites and regimes, to determine the optimal eDNA soil sample depth, volume, and numbers per unit area, similar to the work of Valckx et al. (2011) for hand-sorting and chemical extraction. The collection of soil samples for eDNA analysis is less time and labour intensive in the field than hand-sorting, but this must be traded off against more time spent in the lab preparing the samples for sequencing and subsequent bioinformatics. Future streamlining could further reduce the sampling effort needed, including the use of faster decontamination techniques in the field (for example Foucher et al., 2020).

Achieving a reliable indicator of species abundances is a key challenge that remains for soil eDNA sampling (Kestel et al., 2022). As well as diversity, earthworm abundance is very important for agricultural soil health as it affects soil functioning, how quickly earthworms can improve degraded soils and the rate at which they spread to new areas (Bertrand et al., 2015; Capowiez et al., 2014; Mathieu et al., 2010; Schon et al., 2017). Whilst the eDNA approach shows strong correlations between the relative abundance of sequences and percentage of adult earthworms ascribed to each species identified at the scale of the pooled samples across all locations and field treatments, inferring species abundance from the eDNA sequence numbers is problematic. Raw sequence numbers can be distorted during the PCR amplification, pooling and sequencing processes (Pinto and Raskin, 2012; Kebschull and Zador, 2015; Fonseca, 2018). A common solution in metabarcoding studies is to use relative abundance data instead, but this is also not without criticism (Pinto and Raskin, 2012; Lovell et al., 2015; Jian et al., 2020). Our results suggest that caution should be taken when trying to infer actual abundance patterns from soil eDNA sequence numbers and the relative abundance data they generate at the sample level, due to high within-sample variability between the eDNA measures and hand-sorting abundances (Figure S5). Sample occupancy proportion has been suggested to be a suitable alternative proxy for abundance in eDNA studies (Hänfling et al., 2016), and this shows a strong correlation with the hand-sorting abundances. However, this only gives an overall abundance estimate for each species across the whole sample area and does not give any information on abundances within and between samples. Progress is being made in developing solutions that will allow more rigorous estimation of abundances from eDNA sampling, including the use of new statistical techniques and quantitative PCR (Lovell et al., 2015; Spear et al., 2021; Yates et al., 2019).

The results from all methodologies clearly show a positive increase in earthworm species diversity in the samples that had been converted from arable to ley management. It is interesting that, despite the higher species richness in the ley strips, the communities were not found to be distinct when analysed through NMDS and PERMANOVA. The earthworm populations in the ley strips must have been seeded by the arable field populations (as the strips were directly sown within the arable fields) and there was no evidence of recruitment from the field margins or hedgerows between ley strips that were continuous to the field edge or separated by a barrier and fallow (Prendergast-Miller et al 2021). Furthermore, the conversion to ley may have been too recent (33 months) to show significant divergence between the communities. In addition, some movement of earthworms between the arable and ley cannot be ruled out, given the estimates of active dispersal rates reported in the literature for the species found here (for example Eijsackers, 2011; Butt et al., 2004; Marinissen and van den Bosch, 1992).

In addition to evaluating the earthworm eDNA methodology, our soil pit biomass and hand sorting recording of earthworms in the arable fields and the ley strips in their third growing season have, importantly, revealed that increasing the duration of grass-clover leys reintroduced into long-term intensively cultivated arable fields to 3 years substantially enhances the recovery of earthworm populations over those seen by Prendergast-Miller et al. (2021) after 2 years. The degree of increase in earthworm population densities in the clover-rich leys by the third year, considerably surpassing those in grassy field margins and even permanent grasslands reported for adjacent fields, was unexpected, but may be explained by the previously noted large stimulation of earthworm abundance by clovers introduced into grasslands, presumably providing a better-quality food-source (Van Eekeren et al., 2009).

The rapid reproduction of the earthworms in the leys, as indicated by the abundance of juveniles, and their continuing rapid population growth, dominated by the common endogeic earthworm *Al. chlorotica*, coincided with substantial improvements in soil aggregation, carbon sequestration into soil macroaggregates and soil hydrological functioning, as documented in parallel studies in the SoilBioHedge project (Hallam et al., 2020; Berdeni et al., 2021; Guest et al., 2022). Some of these improvements in the leys were directly attributed to the effects of earthworms by comparing the experimental depletion and enrichment of earthworm populations in field-based mesocosms (Hallam et al. 2020). This included increases in hydraulic conductivity through pores, in plant-available water, and in water-holding capacity, and macropores made a greater contribution to water flows (Hallam et al., 2020). It therefore seems likely that the impressive annual increases in earthworm abundance in the leys over three years was driven by positive feedbacks, whereby the actions of the earthworms in improving soil structure and functioning (Berdeni et al., 2021; Guest et al., 2022), and the minimal disturbance of the periodically mown evergreen grass-clover ground cover, provided a favourable environment for earthworm reproduction and soil health improvement. Our finding that the earthworm population sustained an increase of over 300 individuals m^-2^ from the second to third year of the ley has policy relevance for UK agriculture, justifying the incentivising of keeping leys in arable rotations for 3 years to further improve soil health.

## Conclusion

Environmental DNA sampling represents a promising alternative to the conventional ways of measuring earthworm diversity in agroecosystems that could help to increase standardisation, reduce time spent in the field and tackle some of the existing sampling biases. Our results show that eDNA can be used to sample active agricultural fields and detect fine-scale changes in earthworm diversity brought about by different management practices, making it a good candidate for use in wider soil health monitoring programmes. Future research could help to optimise and refine the technique further, by addressing some of the key unanswered questions surrounding eDNA deposition and degradation in soils, the contribution of legacy eDNA and obtaining reliable abundance information from sequence data. Based on the results of this study and the previous work described here, it is anticipated that eDNA sampling offers the potential to be an important tool for monitoring earthworm diversity and soil health in agriculture, just as it has become a vital instrument for monitoring marine and aquatic habitats. In addition to the insights into earthworm eDNA sampling, this study also builds upon the evidence base showing the value of converting arable soils to grass-clover ley. Our study shows that significant increases in earthworm populations continue to increase significantly into the third year of ley management, so adopting grass-clover leys for longer durations could provide substantial soil health benefits in agroecosystems.

## Acknowledgements

The study was supported by a University of Sheffield Grantham Centre for Sustainable Futures studentship to JL, Natural Environment Research Council (NERC) grant NE/M017044/1 to JRL and a NERC Environmental Omics Facility (NEOF) Award to PJW.

## Author contributions

Conceptualization, J.L., P.J.W., J.R.L, T.B. and H.H.; Formal analysis, J.L., K.H.M., H.H and J.R.L.; Funding acquisition, J.L., P.J.W., and J.R.L.; Investigation, J.L., M.L., G.H. and K.H.M.; Methodology, J.L., P.J.W., J.R.L., T.B., G.H., K.H.M. and H.H; Resources, T.B.; G.H., K.H.M. and H.H.; Software, H.H. and K.H.M.; Supervision, P.J.W, J.R.L and T.B.; Visualisation, J.L. and J.R.L.; Writing – original draft, J.L.; Writing – review and editing, J.L., P.J.W, J.R.L and T.B.

