## Supplementary Materials for "Environmental DNA is more effective than hand sorting in evaluating earthworm biodiversity recovery under regenerative agriculture"

### 1. Supplementary Methods

#### 1.1 Protocol for the extraction of eDNA from soil samples using a phosphate buffer

This protocol has been adapted from a protocol outlined in Taberlet *et al.*, 2018: Environmental DNA for Biodiversity Research and Monitoring (page 37).

I chose to run through the below protocol using three soil samples, taking two replicates of each (i.e. six 15 g soil samples) at a time, plus a negative control. This was due to the falcon rotator being used only having six slots, but also as taking on more samples at a time would have led to more difficulties during the time-sensitive steps. I usually ran through the protocol twice a day, getting through six soil samples (12 replicates) out of 96 total soil samples – plus two negative controls – per day. However, this can be increased or decreased as needed. All of the steps were performed in a lamina-flow hood, unless stated. The protocol uses equipment and chemicals from the Macherey-Nagel NucleoSpin® Soil Kit.

1. Ensure all equipment has been properly cleaned and sterilised before use. Cleaning with 10% bleach after each use and placing in the UV lamina-flow hood overnight is best.
2. Remove the soil samples (in sealed bags) from the -20 °C freezer and defrost in the store room overnight, for use the next day.
3. Prepare 500 ml of saturated phosphate buffer solution by adding 0.985 g $NaH_2PO_4$ and 7.35 g $Na_2HPO_4$ to 500 ml $ddH_2O$. Mix thoroughly until phosphate is dissolved, then autoclave. This buffer cannot be kept for longer than 24 hours after autoclaving, so new buffer will need to be made fresh regularly.
4. Label 50 ml falcon tubes with unique codes corresponding to the soil sample they will correspond to, or with a negative control number.
5. Homogenise the collected soil in the sealed bags, by working the soil with your fingers – taking care that stones do not pierce the bag itself. Often, particularly with the arable soils, this will lead to the soil coalescing as one large clump. When this happens, continue to work the clump like a ball of clay or dough, to ensure proper homogenisation of the soil.
6. Weigh out 15 g of soil from each bag using a disposable plastic spoon and wooden toothpick. Try to sample different parts of the soil in the bag to make up the 15 g (i.e. not just one single large lump), and if the soil is already in a single clump, take pieces from various parts of it. Add this soil to the relevant 50 ml falcon tube and seal.

7. To get two replicates from each bag (i.e. two 15 g samples), repeat step 6. Then dispose of the used plastic spoon, toothpick and weighing boat between each sample bag.
8. Clean and sterilise the weighing scales and surrounding areas by wiping down with 10% bleach between each soil sample bag, and change your gloves. I found that laying a paper towel down next to the scales and where homogenisation took place was useful, to catch any dropped soil and make disposing/subsequent cleaning easier.
9. Add 15 ml of the autoclaved phosphate buffer to each of the falcon tubes containing soil. Include a negative extraction control (a falcon tube that contains only phosphate buffer). If the phosphate buffer has come fresh from the autoclave and is still hot, run the exterior of the bottle under the cold tap until it cools.
10. Use the falcon rotator to rotate the falcon tubes for 15 minutes (out of the lamina-flow hood).
11. While the tubes are rotating, distribute 250 ul of SB buffer into Eppendorf tubes, one tube for each extraction.
12. While the tubes are rotating, put the spin columns on the vacuum manifold ('hedgehog') connectors, close the tops and label.
13. After 15 minutes, take the falcon tubes out of the rotator and take them down to the 50 ml tube centrifuge. Centrifuge the tubes for 5 minutes at 4700 rpm.
14. While the tubes are in the centrifuge, put the elution buffer SE in the oven and set to 80 °C.
15. Remove the tubes from the centrifuge and wipe them with 10% bleach, before taking them back up to the lamina-flow hood. Be careful to walk slowly and try not to mix the supernatant with the floating debris in the tube!
16. Remove 400 ul of the supernatant from the 50 ml falcon tube and transfer it into the Eppendorf tube containing 250 ul SB buffer. Take the supernatant from around 10 mm above the sediment and try to avoid transferring bits of floating debris with the supernatant.
17. Thoroughly mix the supernatant with the buffer using the same filter tip and transfer the 650 ul mix to the relevant spin column.
18. Put the vacuum on (i.e. open the tap on the connector) and the liquid will pass through the column.
19. After all columns have been loaded and all liquid has passed through, break the vacuum for each column.
20. Load 500 ul of SB buffer to each column and put the vacuum on again.
21. Once all liquid has passed through, break the vacuum and load 550 ul of SW1 buffer. Open the taps.

22. Once all the liquid has gone through, break the vacuum and load 750 ul of the SW2 buffer to the columns. Remember to break the vacuum before loading this buffer, so that it can clean the very top parts of the columns.
23. Put the vacuum on again until all the liquid has passed through.
24. Close the columns and transfer each of them to a 2 ml collection tube without cap, and centrifuge for 2 minutes at 11,000 x g to dry the silica membrane
25. If necessary, remove the columns and tap the collection tubes on a dry paper towel to remove the liquid residue inside, ensuring no contamination between tubes. Return each column to its original collection tube after dabbing.
26. Add 680 ul of SW2 buffer, close the columns and then vortex for 2 seconds.
27. Centrifuge the columns for 30 seconds at 11,000 x g
28. Pour out the liquid from each collection tube into a sink or container, and tab the tube on a dry paper towel to remove excess liquid. Ensure no contamination between each tube occurs, and dilute the solution running down the drain by running the taps for a short time.
29. Return the columns to their collection tubes and centrifuge for 2 minutes at 11,000 x g for drying the silica membrane.
30. Put each column on a labelled collection tube with cap and discard the previous collection tube.
31. Collect the elution buffer SE from the oven, which should now be heated to 80°C.
32. Take this back up to the lamina-flow hood (quickly!) and add 100 ul of elution buffer SE to each column.
33. Wait 1 minute at room temperature, then centrifuge for 30 seconds at 11,000 x g.
34. Remove the columns from the collection tubes and store the DNA extract collected in the tubes in the -20 °C freezer. This extract will likely need to be diluted – x5 is best - before use to limit the influence of PCR inhibitors that are coextracted with the DNA.
35. Make sure all equipment to be reused is cleaned with 10x bleach and UV'ed in the lamina-flow hood before next use. To clean the hedgehog, 10% bleach was allowed to soak in the connectors for 15 minutes before opening the taps. The bung was then taken out and the contents emptied into a sink and rinsed with plenty of water.

#### 1.2 Primer selection and in-silico analysis

The primers used in this study are described in Bienert et al. (2012) and amplify short sequences of mitochondrial 16S rDNA. The two primer pairs consisted of primers 'ewD' and 'ewE', which are 17 bp and 21 bp in length respectively and amplify a region of ~70 bp, and primers 'ewB' and 'ewE', which are both 21 bp in length and amplify a region of ~120 bp.

*In-silico* analysis of the primers was performed prior to selection, to check for appropriate primer binding and sufficient variation between earthworm species within the target region. This was performed using MEGA7 and R version 3.5.0 or later (Kumar et al., 2016; R Core Team, 2020), using sequences publicly available on GenBank® (Sayers et al., 2019) covering 24/29 of UK earthworm species (as described in Sherlock, 2012). The remaining five UK earthworm species that did not have sequence data available were considered either rare or very rare and not likely to be found in agricultural soil habitats (Sherlock, 2012).

#### 1.3 Environmental DNA amplification, purification and sequencing

All forward and reverse primers were tailed with Illumina sequencing primers sites (F 5'-3' TCTACACGTTCAGAGTTCTACAGTCCGACGATC; R 5'-3' GTGACTGGAGTTCAGACGTGTGCTCTTCCGATCT).

The PCR1 conditions consisted of an initial denaturation step at 95°C for 15 minutes, followed by 49 cycles of 94°C for 30 seconds, 57°C for 90 seconds and 72°C for 90 seconds, before ending with final step at 72°C for 10 minutes.

After PCR 1, 4 µl of each PCR product was run on a 1% agarose gel and visualised for 45 minutes at 110 volts and imaged under UV light. All PCR1 products were then frozen at -20°C before proceeding to the next stages.

PCR extracts were purified by performing 0.5:1 and 1:1 magnetic bead cleans using AMPure XP beads.

After bead cleaning, a second PCR (PCR2) was performed to add unique Illumina adaptors and indexes to the libraries. PCR2 preparations were done in semi-skirted PCR plates on ice and consisted of making up a total volume of 20 µl with 1 µl each of Fi5 and Ri7 primers (at 2 µM), 8 µl of PCR1 product and 10 µl of Qiagen Multiplex PCR Master Mix. The samples were then incubated at 95°C for 15 minutes, followed by 12 cycles of 98°C for 10 seconds, 65°C for 30 seconds and 72°C for 30 seconds, with a single final step at 72°C for 5 minutes. A subset of samples was then selected and run on the TapeStation, with an observed increase in amplicon size between the pre- and post-PCR2 samples indicating successful addition of the identifying sequences.

After PCR2, initial pooling of the samples was carried out to a concentration of 40 ng/µl. Twenty-four pools (or 'libraries') were created for each primer pair, after which each library underwent a further 1:1 bead clean. Serial dilutions of the libraries were made prior to quantification with qPCR. On ice, 2 µl of the diluted sample pools were mixed with 6 µl of KAPA SYBR® FAST master mix (including primers) and 2 µl of molecular biology grade sterile water, leaving a reaction volume of 10 µl. Negative controls and standards of known concentrations were included in each qPCR run. The qPCR conditions consisted of 5 minutes at 95°C followed by 35 cycles of 30 seconds at 95°C and 45 seconds at 60°C. The qPCR results were used to calculate the concentrations of the libraries (in nM). Further equimolar pooling was then carried out of the libraries (initially to around 150 nM) and combined resulting in one library pool for each primer pair. Vacuum concentration and resuspension in ddH2O were used to get the final volume of these pools down to 20 µl, before both library pools were then gradually diluted with ddH2O and brought down to a final concentration of 4 nM. Running of the pools on the TapeStation revealed a small peak that indicated some lingering primer dimers, so size selection using the BluePippin system was performed according to the manufacturer's instructions. After a further 1:1 bead clean a repeated inspection of product size peaks on the TapeStation showed the primer dimers had been successfully removed. 10 µl of each library pool was then combined and taken for sequencing. Next generation Illumina MiSeq sequencing (using a MiSeq v2 2x 150bp run) was then carried out at Sheffield Children's Hospital.

### 2. Supplementary Results

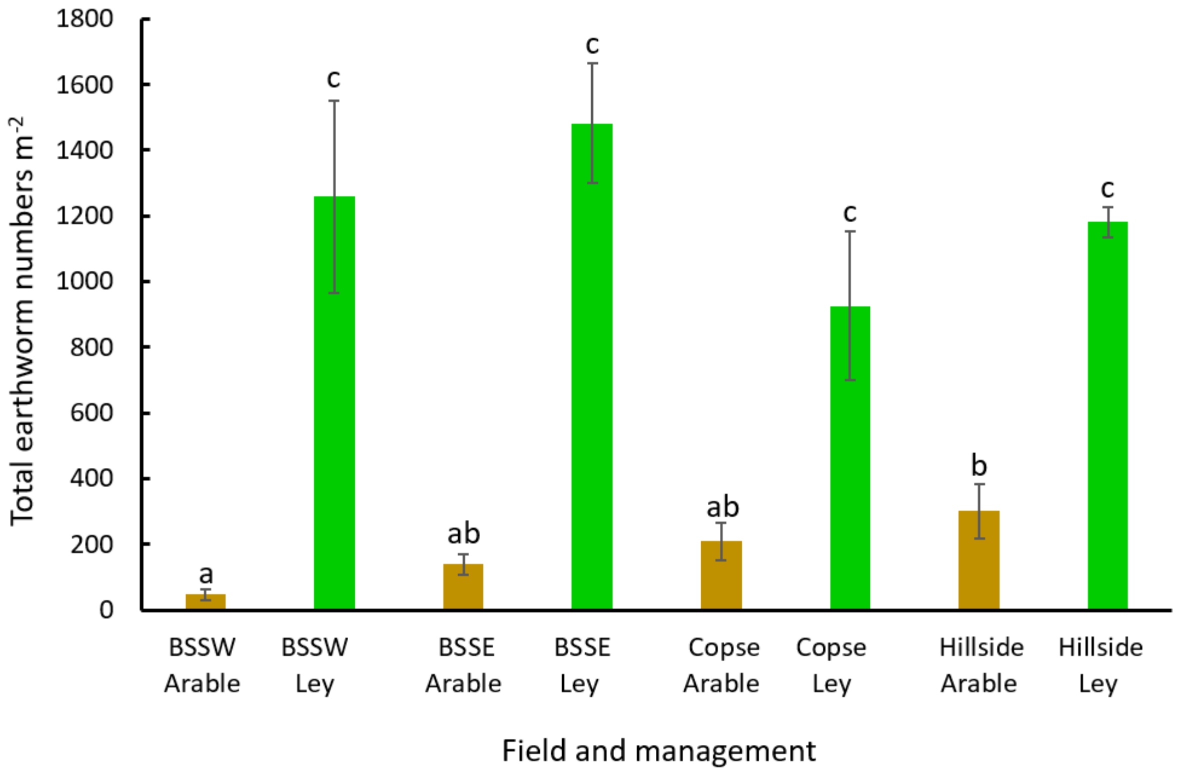


**Figure S1.** Mean earthworm densities per square metre in the arable fields and ley strips, split by field, with standard errors shown. Bars sharing the same letter are not significantly different (Tukey test $p$ >0.05).

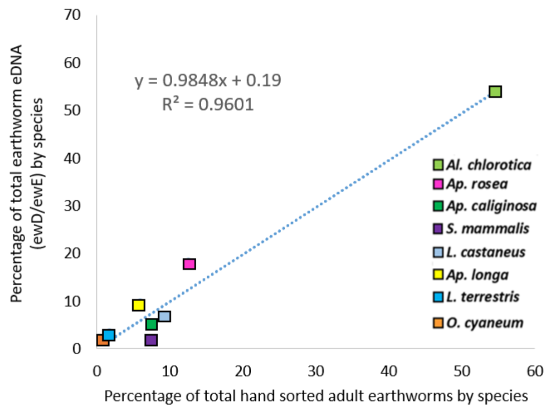


**Figure S2.** The correlation between the percentage of the total earthworm ewD/ewE sequences by species plotted against the percentage of adult earthworms assigned to the 8 species, pooling data from all sampling sites in April 2018. Full species names are given in Table 1.

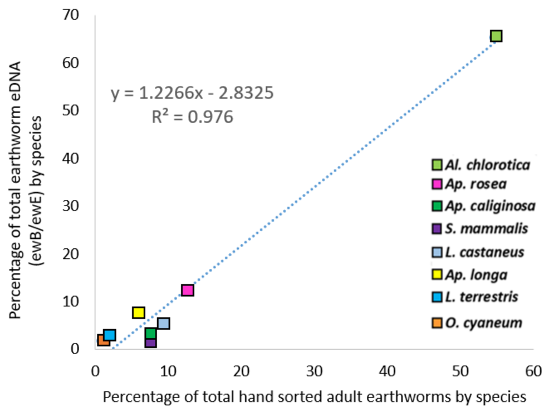


**Figure S3.** The correlation between the percentage of the total earthworm ewB/ewE sequences by species plotted against the percentage of adult earthworms assigned to the 8 species, pooling data from all sampling sites in April 2018. Full species names are given in Table 1.

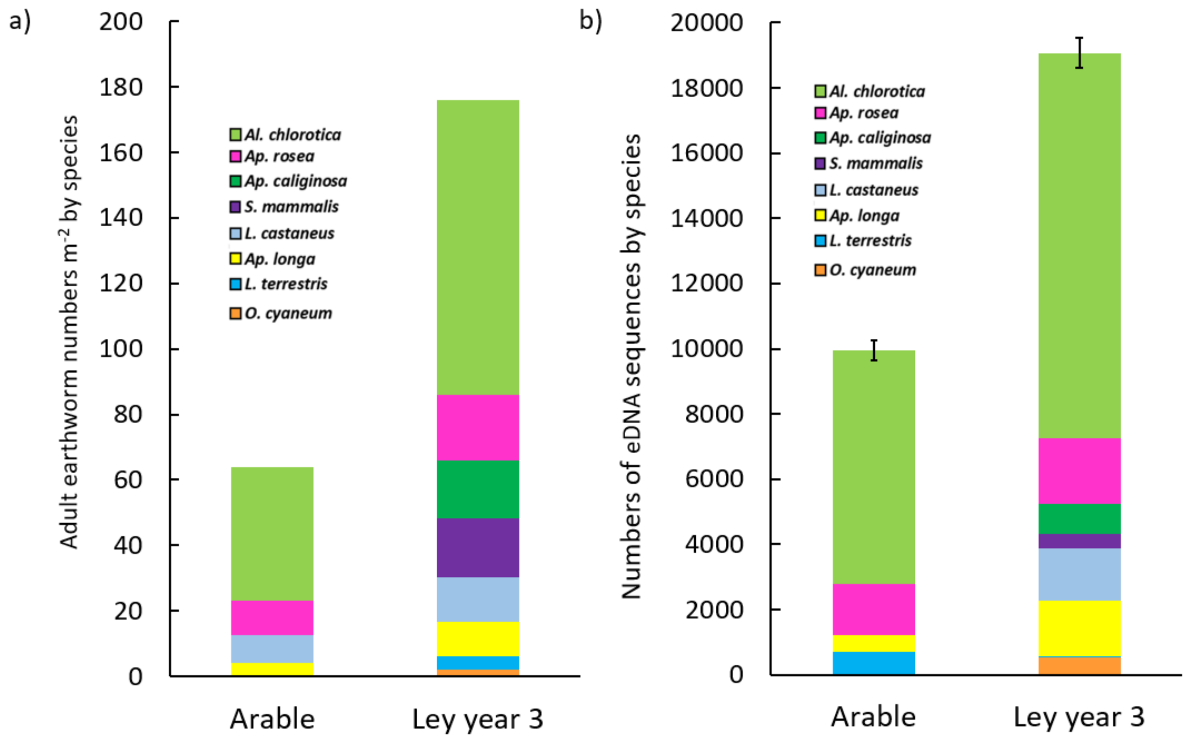


**Figure S4.** (a) Mean adult earthworm numbers per $m^2$, by species for arable and ley parts of the 4 sampled fields and (b) Mean eDNA sequence copy numbers by species using ewD/ewE sequences, for the arable and ley parts of the same 4 fields, with standard errors of the mean total eDNA sequence reads. Full species names are given in Table 1.

a) ewB/ewE – Hand-sorting (Arable)

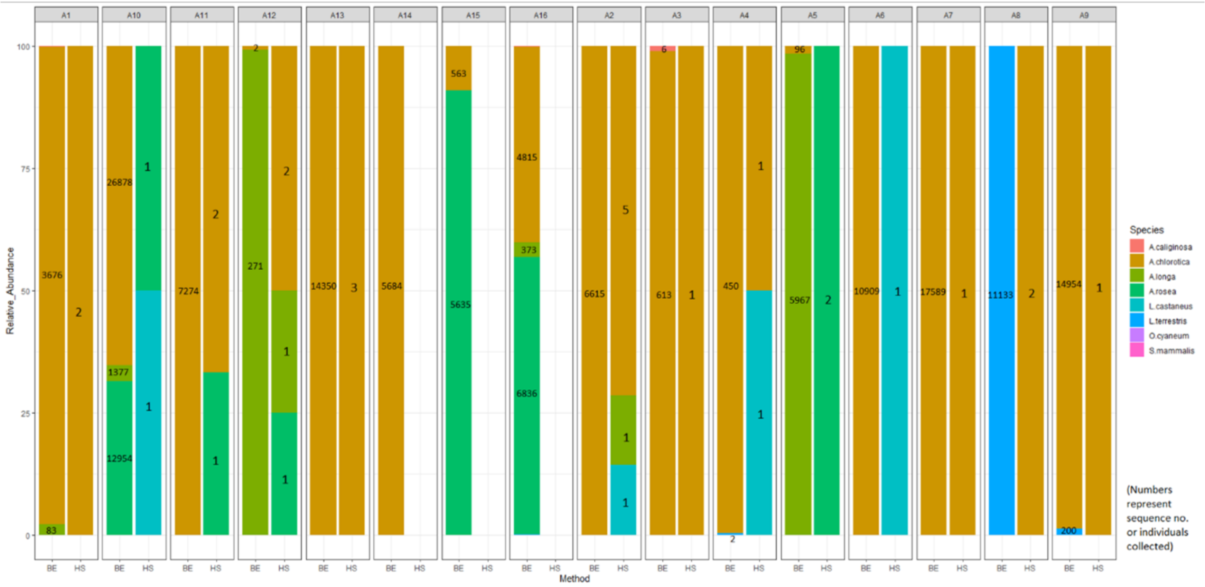


b) ewB/ewE – Hand-sorting (Ley)

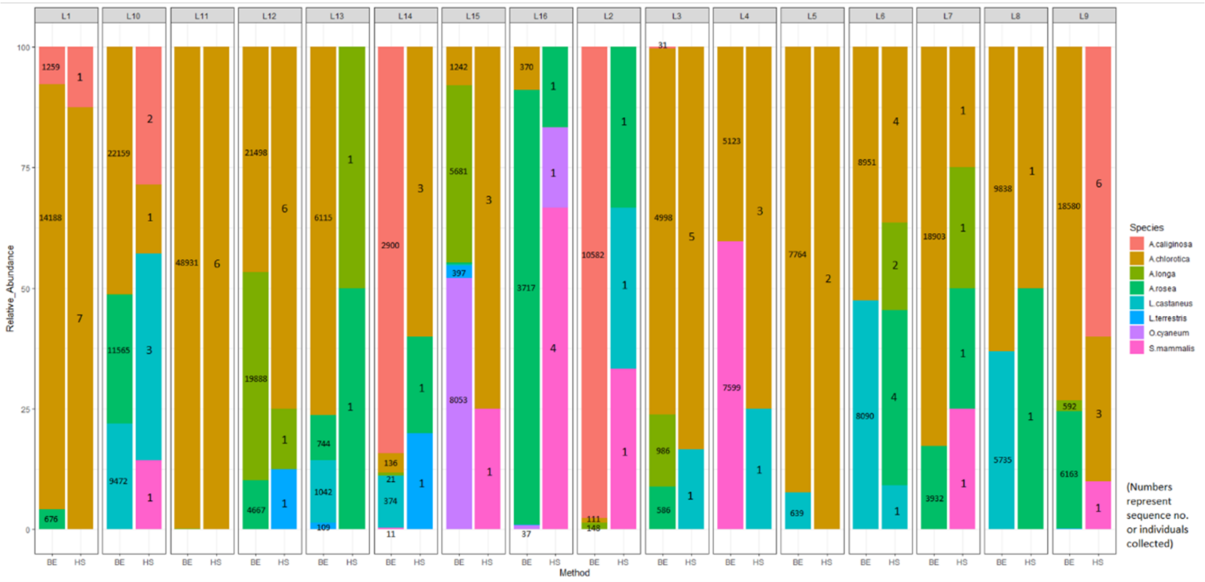


**Figure S5.** Stacked bar charts comparing the relative abundance of species within samples, as measured by the ewB/ewE eDNA method and hand-sorting for the a) arable and b) ley samples. Species relative abundances are marked by the different coloured bars and labelled with the actual sequence numbers and hand-sorting abundances.

#### 2.2 NMDS plots showing difference in earthworm communities across the four fields (1 plot per

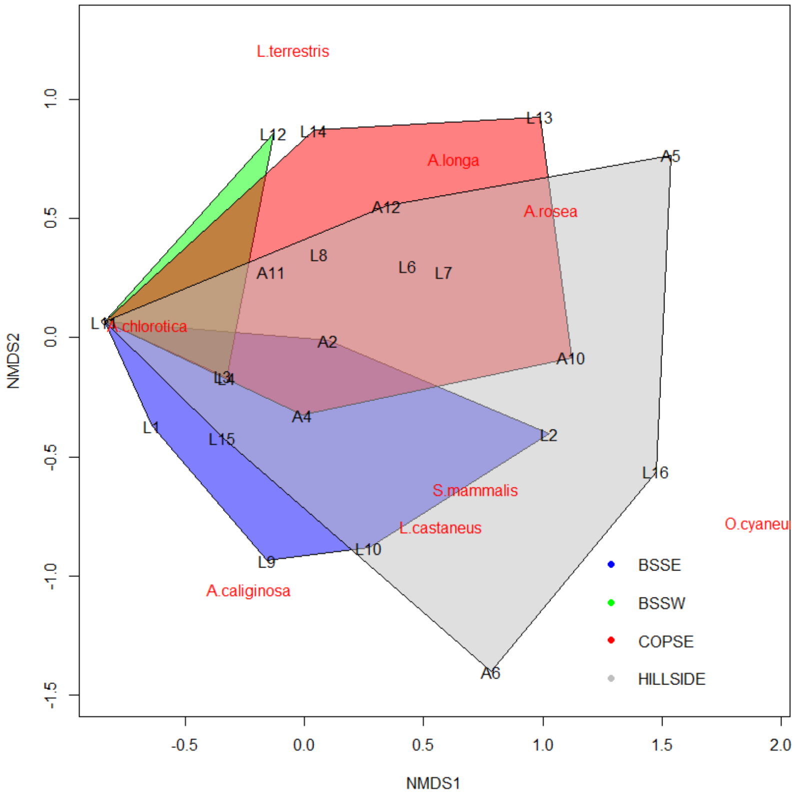


**Figure S6.** NMDS ordination for the results of hand-sorting overlaid by polygons connecting the vertices of points made by the communities in the different fields. The site scores in ordination space are represented by the site labels (arable sites = 'A1', 'A2' etc., ley sites = 'L1', 'L2' etc.), and the species labels are positioned at the weighted average of the site scores.

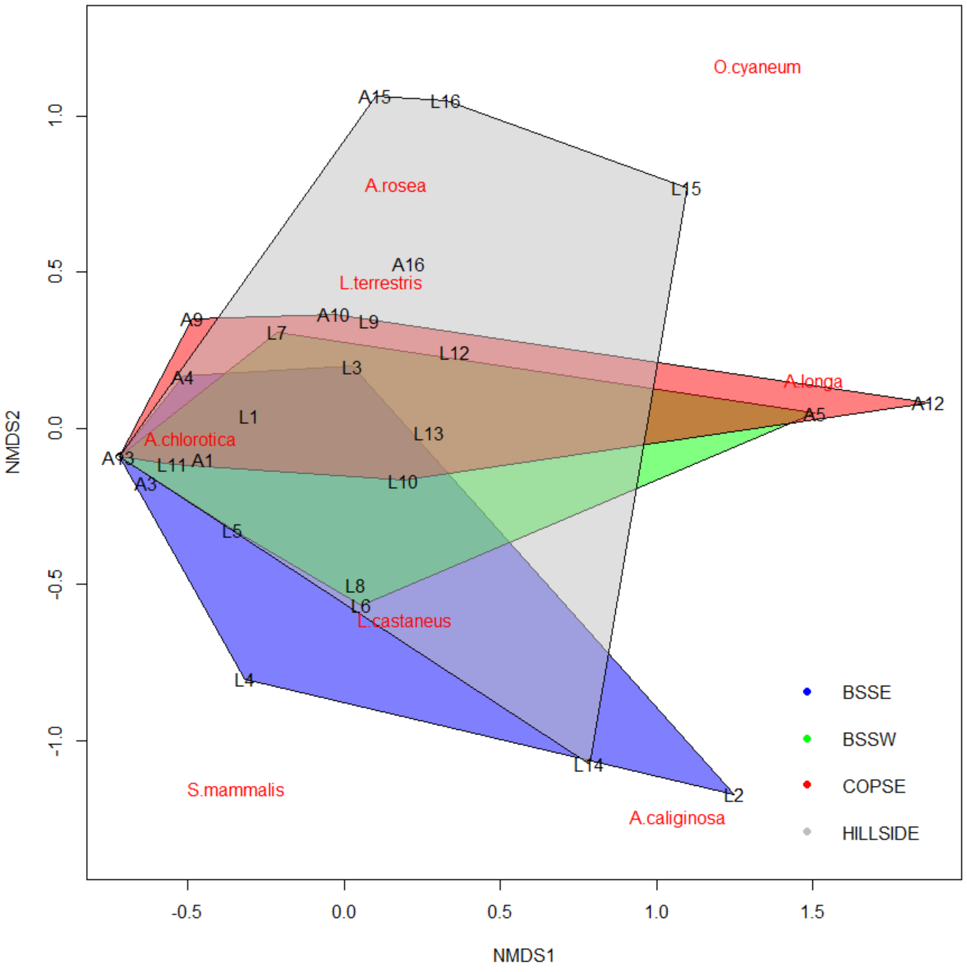


**Figure S7.** NMDS ordination for the results of eDNA sampling using ewB/ewE primers, overlaid by polygons connecting the vertices of points made by the communities in the different fields. The site scores in ordination space are represented by the site labels (arable sites = 'A1', 'A2' etc., ley sites = 'L1', 'L2' etc.), and the species labels are positioned at the weighted average of the site scores.

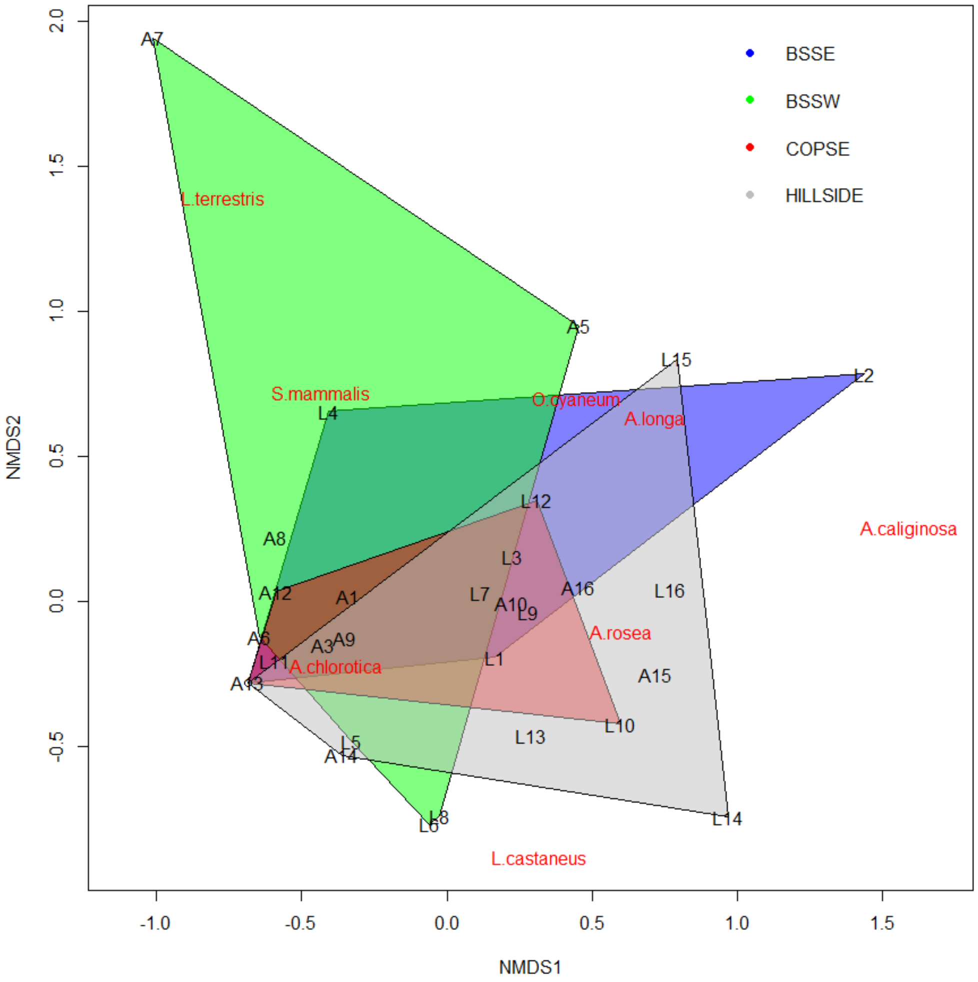


**Figure S8.** NMDS ordination for the results of eDNA sampling using ewD/ewE primers, overlaid by polygons connecting the vertices of points made by the communities in the different fields. The site scores in ordination space are represented by the site labels (arable sites = 'A1', 'A2' etc., ley sites = 'L1', 'L2' etc.), and the species labels are positioned at the weighted average of the site scores.

### 2.3 Correlations of the relative abundance scores for the different sampling methods

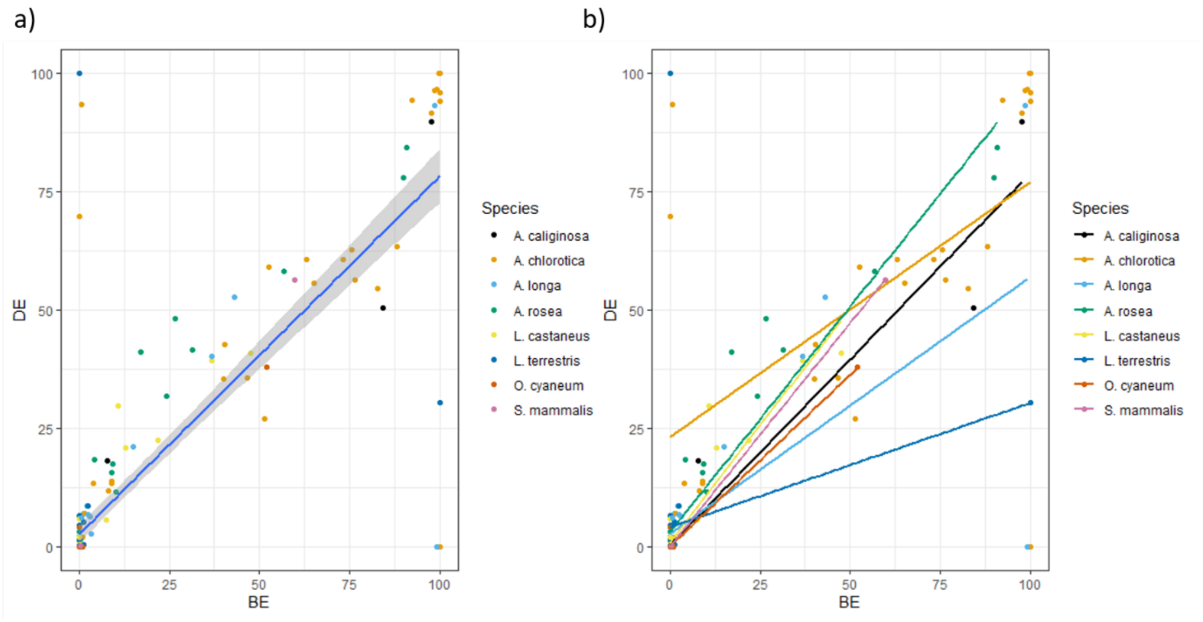


**Figure S9.** Comparing the relative abundances of different species in each sample reported by the ewB/ewE and ewD/ewE primer pairs. A) gives the overall trend with all species grouped together while b) shows the separate trends for each species. 'DE' = ewD/ewE and 'BE' = ewB/ewE.

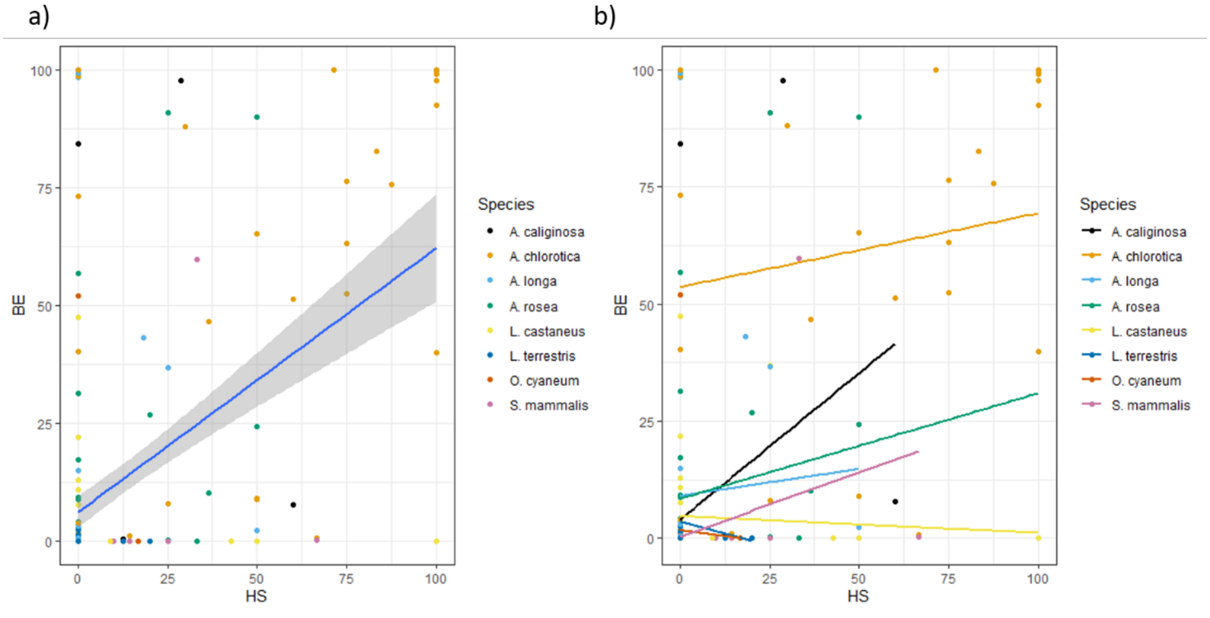


**Figure S10.** Comparing the relative abundances of different species in each sample reported by the ewB/ewE eDNA primer pair and hand-sorting. A) gives the overall trend with all species grouped together while b) shows the separate trends for each species. 'BE' = ewB/ewE and 'HS' = hand-sorting.

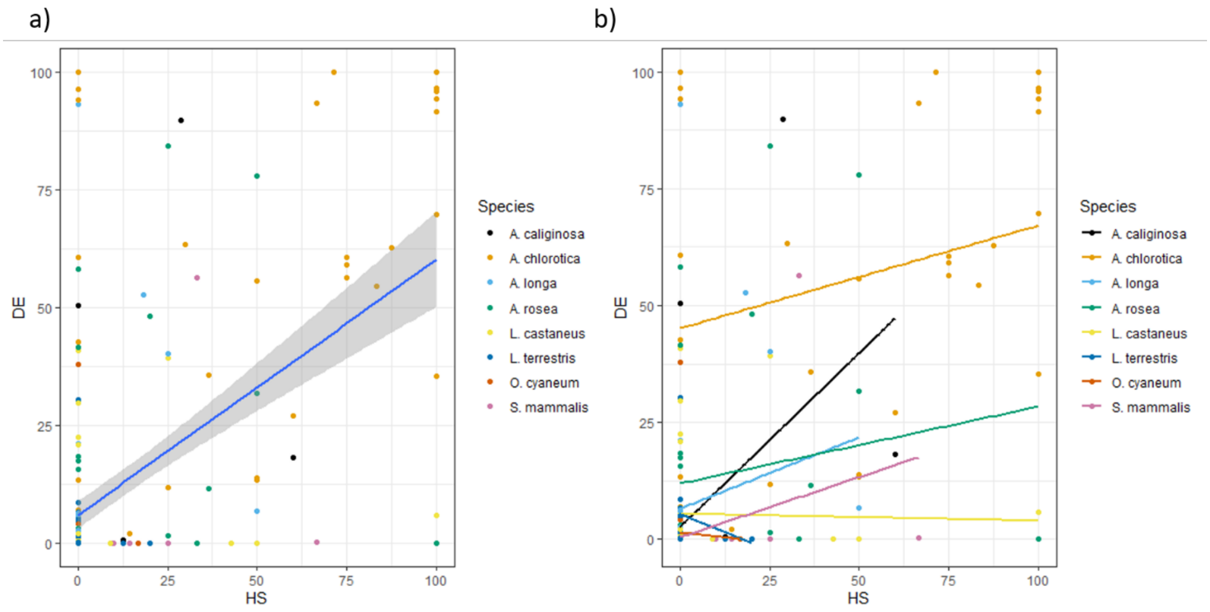


**Figure S11.** Comparing the relative abundances of different species in each sample reported by the ewD/ewE eDNA primer pair and hand-sorting. A) gives the overall trend with all species grouped together while b) shows the separate trends for each species. 'DE' = ewD/ewE and 'HS' = hand-sorting.
